# Neuroimaging Correlates of Emotional Response-Inhibition Discriminate Between Young Depressed Adults With and Without Sub-threshold Bipolar Symptoms

**DOI:** 10.1101/2020.04.25.060111

**Authors:** Jungwon Cha, Sidra Speaker, Bo Hu, Murat Altinay, Parashar Koirala, Harish Karne, Jeffrey Spielberg, Amit Anand

**Author notes:** Corresponding Author: Amit Anand, MD, Center for Behavioral Health, 9500 Euclid Avenue P57, Cleveland, OH 44122.

## Abstract

**Background:** A significant number of subjects with major depression (MDD) exhibit subthreshold mania symptoms (MDD+). This study investigated, for the first time, using emotional inhibition tasks, whether the neurobiology of MDD+ subjects is more akin to bipolar disorder depression (BDD) or to MDD subjects without any subthreshold bipolar symptoms (MDD−).

**Method:** This study included 118 medication-free young adult subjects (16 – 30 yrs.): 20 BDD, 28 MDD+, 41 MDD−, and 29 HC subjects. Participants underwent fMRI during emotional and non-emotional Go/No-go tasks during which they responded for Go stimuli and inhibited response for happy, fear, emotional (happy + fear) and non-emotional (gender) faces No-go stimuli. Linear mixed effects (LME) analysis for group effects and Gaussian Process Classifier (GPC) analyses was conducted.

**Results:** MDD− group compared to both the BDD and MDD+ groups, exhibited significantly lower activation in parietal, temporal and frontal regions (cluster-wise corrected p <0.05) for emotional inhibition conditions vs. non-emotional condition. No significant differences were found between BDD and MDD+ groups. GPC classification of emotional vs non-emotional response-inhibition activation pattern showed good discrimination between BDD and MDD− subjects (AUC: 0.70; balanced accuracy: 70% (p = 0.006)) as well as MDD+ and MDD− subjects (AUC: 0.72; balanced accuracy: 67% (p = 0.015)) but less efficient discrimination between BDD and MDD+ groups (AUC: 0.68; balanced accuracy: 61% (p = 0.091)). Notably, classification of the MDD− group was weighted for left amygdala activation pattern.

**Conclusion:** Using an fMRI emotional Go-Nogo task, MDD− subjects can be discriminated from BDD subjects and MDD+.

## Introduction

It has been estimated that 30-55% of patients presenting with major depression (MDD) have additional sub-threshold hypo(manic) symptoms (MDD+). MDD+ subjects are thought to be at a higher risk for developing bipolar disorder (BD) (Coryell, Endicott et al. 1995, Fiedorowicz, Endicott et al. 2011). However, the neurobiology of MDD+ subjects has not been adequately studied to make a determination whether MDD+ are more akin to BDD or to MDD− subjects. In particular, young subjects presenting with both depression and subthreshold mania (MDD+) present a conundrum for clinicians as the duration of illness is usually too short to make a determination about the stability of the diagnosis. Therefore, there is a critical need to differentiate and identify objective classifiers, which may help in differentiating BDD, MDD− and MDD+ in young adult subjects.

Neuroimaging characteristics for BD may provide such potential classifiers. Neuroimaging studies in BD have implicated abnormal corticolimbic activation and connectivity in BD though no single abnormality has been robustly validated through replication studies (Strakowski, Adler et al. 2012, Phillips and Swartz 2014). fMRI task-induced activation paradigms involving affective faces have frequently been used in studies of mood disorders, including studies comparing MDD and BDD in order to isolate BDD-specific characteristics (Redlich, Almeida et al. 2014, Redlich, Dohm et al. 2015, Han, De Berardis et al. 2019). However, a critical gap remains, given that very few studies have examined this question *within* the MDD group; in other words, comparing MDD+ to MDD subjects who do not have subthreshold mania symptoms (MDD−). Fournier and colleagues reported that greater right-amygdala activity to the happy faces in adults with MDD was associated with higher levels of subthreshold manic symptoms experienced across the lifespan (Fournier, Keener et al. 2013). A recent study of remitted young depressed subjects found a difference in resting state brain connectivity in subjects with subthreshold bipolar symptoms (Kling, Bessette et al. 2018). Thus, there are indications that affect-face-related activation may be used to characterize MDD+ individuals, but this has not been explicitly tested. In the present study, we have attempted to fill this critical gap in the literature by testing whether fMRI activation during response-inhibition to emotional task can differentiate and accurately classify MDD+ and MDD− individuals.

BD is frequently associated with difficulty inhibiting responses, particularly to emotion-laden stimuli. One commonly used task to assay these difficulties is a variant of the Go/No-go task, in which a subject has to inhibit responses to a rare No-go stimulus in the context of speeded responses to a common Go stimulus. Both non-emotional (e.g., letters) and emotional (e.g., faces depicting an emotion) stimuli are commonly used in this paradigm.

A number of studies have used emotional Go/No-go tasks to investigate BD-related differences in activation. For example, BD has been associated with differential activation on emotional Go/No-go tasks in the temporal cortex, OFC, insula, caudate, and the dorsal anterior and posterior cingulate cortices (Wessa, Houenou et al. 2007, Townsend, Bookheimer et al. 2012, Hummer, Hulvershorn et al. 2013).

Although several studies have used close relatives of bipolar subjects to identify high risk endophenotypes (Piguet, Fodoulian et al. 2015, Vierck, Porter et al. 2015, Frangou, Dima et al. 2017), MDD+ subjects have not yet been used to identify clinically applicable activation patterns, with the ultimate goal of applying these patterns in clinical work to identify at-risk individuals.

Typical fMRI studies in this area employ univariate analyses to find group-related differences. Though information gained via these methods provides valuable insights into possible BDD pathophysiology, they have less individual-level-clinical utility for differentiating between groups (Frangou, Dima et al. 2017, Amit Etkin 2019). In comparison, multivariate machine learning techniques, such as support vector machines (SVM) or Gaussian Process Classifiers (GPC), can be to provide individual-level classification accuracy of an fMRI task. For example, Frangou and colleagues used GPC to classify BD and relatives of BD at-risk for developing BD, using a 3-back working memory task (Frangou, Dima et al. 2017). Such individual level analyses could have utility in clinical situations which require correctly e.g. differentiating between MDD+ and MDD− .

The aim of this study was to ascertain the differentiating and classification-accuracy of emotional Go/No-go task induced fMRI activation in MDD+ and MDD− subjects. Based on previously described findings, we hypothesized that during emotional response-inhibition, MDD+ and BDD participants will show a more similar pattern of activation for emotional vs. non-emotional response-inhibition tasks, which will be different from the pattern seen in MDD− subjects. To test this hypothesis, we examined, for the first time, fMRI task induced activation differences between medication-free young subjects with BDD, MDD+, MDD−, and HC as participants completed emotional (happy, fearful) and non-emotional (gender recognition) Go/No-go tasks. Next, we used a GPC machine learning algorithm to evaluate classification performances of fMRI activation patterns and identify brain areas which contributed the most to the classification.

## Materials and Methods

### Subjects

Bipolar and MDD participants ages 15-30 years who were medication-free for at least 2 weeks were recruited from the outpatient psychiatry clinic at the Cleveland Clinic and by advertisement. Healthy control participants were recruited by advertisement to the community. Inclusion and Exclusion criteria are presented in the supplement.

### Depression subgroup ascertainment using best practices

Three psychiatrists independently reviewed all information available for each subject and classified all MDD subjects as MDD+ or MDD− at the conclusion of the study (Koirala, Hu et al. 2019). Based on review of previous studies (Angst, Gamma et al. 2003, Zimmermann, Brückl et al. 2009, Fiedorowicz, Endicott et al. 2011, Merikangas, Jin et al. 2011, Koirala, Hu et al. 2019), which have used varying definitions for subthreshold mania symptoms, we formulated conservative criteria and defined MDD+ as: euphoric mood with at least 2 mania symptoms or increased irritability with 3 mania symptoms if only the latter was present, as well as if full mania symptoms were present then duration to be for less than 4 days. All Other MDD subjects who did not have subthreshold symptoms, no family history of BD, and no history of psychosis (Angst, Gamma et al. 2003, Zimmermann, Brückl et al. 2009, Fiedorowicz, Endicott et al. 2011, Merikangas, Jin et al. 2011, Koirala, Hu et al. 2019), were ascertained to be MDD− . Subsequently, the classification of each of the subjects was discussed between all three psychiatrists and a consensus best-estimate classification was agreed upon. (Nurnberger, McInnis et al. 2011).

Naturalistic follow-up for change in diagnosis: as an exploratory aim of the study, MDD subjects who wanted to be treated were started on a selective serotonin reuptake inhibitor (SSRI) and followed up for up to 2 years. The course of the larger sample, which was clinically followed up has been reported previously (Koirala, Hu et al. 2019). Any change in diagnosis to BD was recorded.

### Procedures

#### Tasks

We used a modified emotion Go/No-go fMRI paradigm similar to that used in other studies (Wessa, Houenou et al. 2007, Hummer, Hulvershorn et al. 2013) with fearful, happy and neutral facial stimuli showing both male and female faces as depicted and described in Supplemental Figure S-1. Emotional inhibition task conditions consisted of emotional No-go blocks in which (Happy or Fearful) No-go stimuli were interspersed among Neutral Go stimuli. For emotional inhibition regardless of valence, Emotional Inhibition (Happy No-go + Fearful No-go) condition was created. For emotional inhibition tasks a contrast with each of the emotional No-go conditions with the neutral Go condition was created. Neutral Go rather than emotional Go was used as the control stimulus for all emotional No-go blocks to avoid responses to strong repetitive emotional stimulus dominating the activation response. No-go stimuli were interspersed among Neutral (Female faces and Male faces) Go stimuli (Supplemental Figure S-1). Non-emotional inhibition condition was constructed consisting of (Female + Male) No-go blocks and Go blocks.

#### Behavioral Analysis

To examine performance differences between groups, Go and No-go accuracies and reaction times were compared using SPSS (Version 21, IBM, Chicago, Illinois) via one-way analysis of variance (ANOVA) tests with post hoc t-test and Bonferroni correction for multiple comparisons as shown in Supplemental Table 1.

#### Imaging procedure and preprocessing: details are presented in online supplement Functional neuroimaging data analysis

A General Linear Model was implemented with a delayed boxcar waveform to model blood oxygen level dependent (BOLD) signal changes in relation to conditions. To deal with the arbitrariness of BOLD signal varying across brain regions and subjects, the data was scaled at the first level (within-subject) (Chen, Taylor et al. 2017). Six movement parameters, calculated during the realignment, were used as covariates of no interest to account for variations in signal from movement artifacts. First level analyses produced beta images for each condition of interest (happy no-go, fearful no-go, neutral go, male no-go, female no-go, male go, and female go) from each participant using 3dDeconvolve from AFNI.

A mask was created of average effect of each task condition across all subjects at voxel-wise threshold p < .001 (uncorrected) which would represent the cluster-wise corrected significance of p < .05 to be used for second level group effect analysis for each condition respectively. To calculate the cluster-wise corrected significance threshold for a given voxelwise threshold p-value, we used 3dClustsim’s non-Gaussian auto-correlation function (ACF) which uses the smoothness parameters estimated by the 3dFWHMx function from AFNI (Forman, Cohen et al. 1995, Cox, Chen et al. 2017).

We conducted second level linear mixed-effects (LME) analysis with a whole brain mask using 3dLME function in AFNI (Chen, Saad et al. 2013). F-statistics for main effect of group × condition was calculated for these contrasts using 3dLME. We used group as a between-subject factor and condition as a within-subject factor, controlling for task accuracy and reaction time with a within-subject quantitative variable, and age, race, and scanner type with between-subject covariates to identify the effect of group on the pre-specified contrasts.

Additionally, we conducted main effects of group × condition analyses of Go only blocks contrasted with Rest condition, to ensure that Go blocks were not driving significant results in No-go – Go contrasts. Significant Go/No-go results are only reported only if analogous Go vs. Rest clusters were not independently significant.

For differences across patient groups, we performed a second level linear mixed-effects (LME) analysis for four groups using age, race, scanner type, reaction time and task accuracy as covariates to exclude the effect of these confounding variables.

Main effect of group × condition was examined at voxel-wise threshold p < .001 (uncorrected) threshold and cluster-wise corrected significance of p < .01 using the average effect of condition mask. We used more stringent cluster-wise corrected threshold of p-value to include the most significant clusters. The location of the cluster was identified using the location of the peak voxel using the MNI Atlas. Cluster sizes required for corrected significance are given below under Results.

Subject-specific beta coefficients from each main effect cluster were analyzed in SPSS (Version 21, IBM, Chicago, Illinois) for post-hoc pair-wise group significant differences using Bonferroni correction for multiple comparisons (Table 2).

#### Three patient group analysis

The MDD− group had a lower mean HAM-D score than the MDD+ and BDD groups. Therefore, to examine for any effect of differences in HAM-D depression scores on differences seen between the three patient groups results seen in the four group analysis, we also conducted a three group analysis for the patient groups while excluding the HC group. In this three group analysis, HAM-D score was additionally used as a covariate beside the other covariates included in the four group analyses. YMRS scores were low in all depressed patient groups but were significantly different in the BDD and MDD+ groups compared to the MDD− group. However, YMRS scores were not used as covariates as they were the inherent characteristics of the MDD+ and the BDD groups. The results of three group analysis were essentially the same as the four group analysis in terms of brain regions and post-hoc differences between the patient groups (Supplementary Table 3). The four-group analysis is presented below to also include healthy subjects in the ANOVA model.

### Multivariate pattern classification

Binary Gaussian Process Classifier (GPC) was performed in the Pattern Recognition for Neuroimaging Toolbox (PRoNTo) (www.mlnl.cs.ucl.ac.uk/pronto/) using contrasts based on whole-brain beta images for each condition of interest (happy No-go, fearful No-go, emotional No-go, neutral go, male No-go, female No-go, non-emotional No-go, male go, and female go).

GPC is a supervised learning method for classification, which builds a model with (Gaussian) predictive probabilities of class membership, which allows us to estimate uncertainty in the prediction (Rasmussen 2003). Age, race, scanner type and 17-item Hamilton Depression (HAM-D) were used as covariates in the classifiers. Each classifier used the mask computed by the average effect of each condition across all subjects in the corresponding group-pair at voxel-wise threshold of p < .001 (uncorrected), cluster-wise corrected significance of p < .05. Each classifier was trained using a leave-one-out per group cross validation (CV). In order to perform the CV, one subject in each group was selected and then allocated to the test set. The primary metric for evaluation of classification performance was the area under the receiver operating characteristics (ROC) curve (AUC). Moreover, due to an imbalanced dataset, standard accuracy may not be a valid way to examine the accuracy of the classifier. In contrast, sensitivity (true positive rate) and specificity (true negative rate) better accommodate imbalanced data. Thus we calculated balanced accuracy (average of sensitivity and specificity) which is thought to be a more valid metric for interpreting the classification for an imbalanced dataset (Brodersen, Haiss et al. 2011). A threshold of 0.5 for class-labels was used. Next, permutation testing was performed for each classifier with 1000 permutations, which provided a p-value for the balanced accuracy of classification (Nichols and Holmes 2002, Golland and Fischl 2003, Noirhomme, Lesenfants et al. 2014).

Finally, we computed whole-brain discrimination maps for each classifier to visualize the quantitative contribution of each voxel to the classifier’s decision.

Classification between pairs of diagnostic groups BDD vs MDD−, BDD vs MDD+, and MDD+ vs MDD− was analyzed. We also looked at classification between (BDD + MDD+) vs MDD− group and the BDD vs (MDD− + MDD+). This was done to investigate whether addition of MDD+ to BDD or to MDD− led to better discrimination as an indirect indicator of whether the MDD+ group was more similar to the BDD group or the MDD− group.

## Results

### Demographics

One-hundred seventy-seven subjects were enrolled in the study. Thirty participants were excluded due to excessive motion or problematic quality during image acquisition. Five participants were excluded because they only had a family history of BD (4 subjects) or psychosis (1 subject) only but no subthreshold symptoms. These subjects were excluded to reduce heterogeneity in the MDD+ sample. Eight HCs were also excluded in order to match race across groups. Two participants from the HC group later on were discovered to have had family history of mental illness and were excluded. One BDD participant was excluded who was judged to give unreliable information. Three participants were excluded since they were on hormones and transition to opposite gender. The final analyses included 118 medication-free subjects: 20 BDD (8 bipolar I and 12 bipolar II), 28 MDD+, 41 MDD−, and 29 HC subjects. The BDD subtypes were combined as a whole and compared with the MDD+ and MDD− groups as there were not a sufficient number of subjects to conduct analysis taking account the BD I and II subtypes.

Demographic Characteristics of each group are presented in Table 1.

**Table 1.** Demographics and Illness Characteristics.

| **Demographics four group ANOVA** |  | **BDD (N=20)** | **MDD+ (N=28)** | **MDD- (N=41)** | **HC (N=29)** | **3-**<br>**Group**<br>**p-value** | **4-**<br>**Group**<br>**p-value** |
| --- | --- | --- | --- | --- | --- | --- | --- |
| Age (years)(mean (SD)) |  | 24.8(4.1) | 23.1(3.5) | 25.1(3.8) | 24.8(3.4) | 0.083 | 0.121 |
| Female (n (%)) |  | 14(70%) | 24(86%) | 27(66%) | 18(62%) | 0.178 | 0.211 |
| Race | Caucasian (n (%)) | 17(85%) | 21(75%) | 39(95%) | 26(90%) | 0.054 | 0.097 |
|  | African American (n (%)) | 3(15%) | 7(25%) | 2(5%) | 3(10%) |  |  |
| Scanner | Scanner 1 (n (%)) | 4(20%) | 16(57%) | 11(27%) | 7(24%) | 0.010 | 0.013 |
|  | Scanner 2 (n (%)) | 16(80%) | 12(43%) | 30(73%) | 22(76%) |  |  |
| Handedness (n of right handed (%)) |  | 18(90%) | 25(86%) | 38(93%) | 22(88) ¶ | 0.641 | 0.818 |
| **Illness characteristics three group ANOVA** |  |  |  |  |  |  |  |
| HAM-D 17 item (mean (SD)) |  | 18.6(5.1) | 19.7(3.7) | 16.6(3) |  | 0.004 |  |
| YMRS (mean (SD)) |  | 2.1(2.5) | 2.5(2.8) | 0.8(1.2) |  | 0.002 |  |
| Age at First Episode (years)(mean (SD)) |  | 13.3(3.7) | 14.7(4.3) | 15.3(4.4) |  | 0.201 |  |
| Duration of illness (years)(mean (SD)) |  | 10.5(5.6) | 8.4(5.6) | 9.8(5.4) |  | 0.390 |  |
| Medication Free Period (weeks)(mean (SD))* |  | 108.5(108.9) | 70(71.6) | 95.7(118.3) |  | 0.551 |  |
| **Classification Characteristics** |  |  |  |  |  |  |  |
| Bipolar I(n) : Bipolar II(n) |  | 8:12 |  |  |  |  |  |
| First degree family history (n (%)) |  | 6(30%) | 4(14%) | 0(0) |  |  |  |
| Psychosis (n (%)) |  | 2(10%) | 3(11%) | 0(0) |  |  |  |

- ¶ 4 subjects are missing information. - \* There were no significant differences in the distribution of medication naïve participants across groups. - ANOVA test was performed on the continuous variables and chi-squared test was performed on the categorical variables. - BDD : bipolar depressed group - MDD+ : major depressive disorder patients with subthreshold mania symptoms - MDD- : major depressive disorder patients who no subthreshold mania symptoms, history of psychosis or family history of bipolar disorder - HC : Healthy Control

**Table 2.**
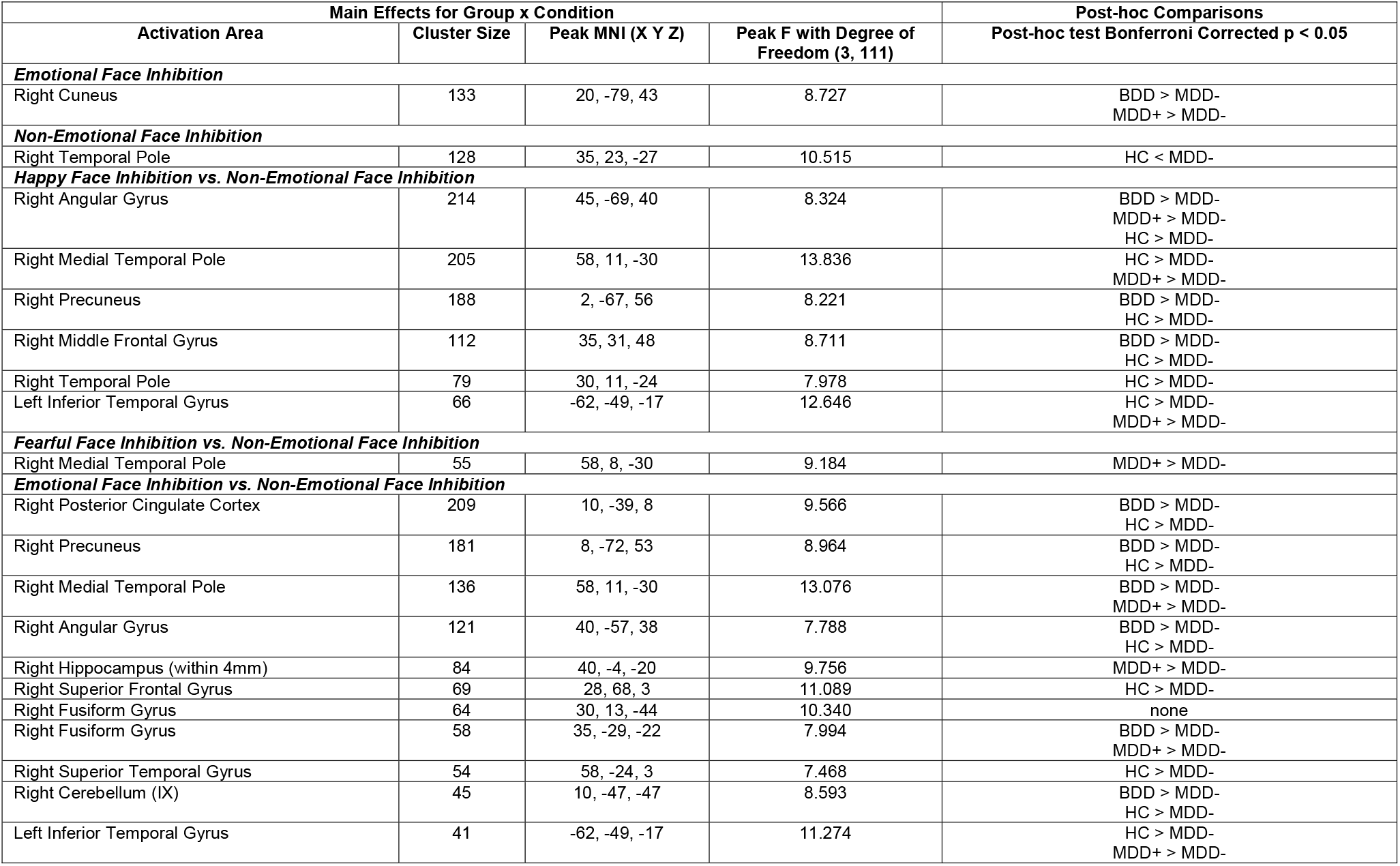
Significant Results of Main Effects for Group × Condition and Post-hoc Comparisons with Bonferroni Correction for Multiple Comparisons.

| Main Effects for Group x Condition |  |  |  | Post-hoc Comparisons |
| --- | --- | --- | --- | --- |
| Activation Area | Cluster Size | Peak MNI (X Y Z) | Peak F with Degree of Freedom (3, 111) | Post-hoc test Bonferroni Corrected p < 0.05 |
| ***Emotional Face Inhibition*** |  |  |  |  |
| Right Cuneus | 133 | 20, -79, 43 | 8.727 | BDD > MDD-<br>MDD+ > MDD- |
| ***Non-Emotional Face Inhibition*** |  |  |  |  |
| Right Temporal Pole | 128 | 35, 23, -27 | 10.515 | HC < MDD- |
| ***Happy Face Inhibition vs. Non-Emotional Face Inhibition*** |  |  |  |  |
| Right Angular Gyrus | 214 | 45, -69, 40 | 8.324 | BDD > MDD-<br>MDD+ > MDD-<br>HC > MDD- |
| Right Medial Temporal Pole | 205 | 58, 11, -30 | 13.836 | HC > MDD-<br>MDD+ > MDD- |
| Right Precuneus | 188 | 2, -67, 56 | 8.221 | BDD > MDD-<br>HC > MDD- |
| Right Middle Frontal Gyrus | 112 | 35, 31, 48 | 8.711 | BDD > MDD-<br>HC > MDD- |
| Right Temporal Pole | 79 | 30, 11, -24 | 7.978 | HC > MDD- |
| Left Inferior Temporal Gyrus | 66 | -62, -49, -17 | 12.646 | HC > MDD-<br>MDD+ > MDD- |
| ***Fearful Face Inhibition vs. Non-Emotional Face Inhibition*** |  |  |  |  |
| Right Medial Temporal Pole | 55 | 58, 8, -30 | 9.184 | MDD+ > MDD- |
| ***Emotional Face Inhibition vs. Non-Emotional Face Inhibition*** |  |  |  |  |
| Right Posterior Cingulate Cortex | 209 | 10, -39, 8 | 9.566 | BDD > MDD-<br>HC > MDD- |
| Right Precuneus | 181 | 8, -72, 53 | 8.964 | BDD > MDD-<br>HC > MDD- |
| Right Medial Temporal Pole | 136 | 58, 11, -30 | 13.076 | BDD > MDD-<br>MDD+ > MDD- |
| Right Angular Gyrus | 121 | 40, -57, 38 | 7.788 | BDD > MDD-<br>HC > MDD- |
| Right Hippocampus (within 4mm) | 84 | 40, -4, -20 | 9.756 | MDD+ > MDD- |
| Right Superior Frontal Gyrus | 69 | 28, 68, 3 | 11.089 | HC > MDD- |
| Right Fusiform Gyrus | 64 | 30, 13, -44 | 10.340 | none |
| Right Fusiform Gyrus | 58 | 35, -29, -22 | 7.994 | BDD > MDD-<br>MDD+ > MDD- |
| Right Superior Temporal Gyrus | 54 | 58, -24, 3 | 7.468 | HC > MDD- |
| Right Cerebellum (IX) | 45 | 10, -47, -47 | 8.593 | BDD > MDD-<br>HC > MDD- |
| Left Inferior Temporal Gyrus | 41 | -62, -49, -17 | 11.274 | HC > MDD-<br>MDD+ > MDD- |

### Task Performance

After correction for multiple pair-wise group differences no significant differences were seen between groups for any of the Go or No-Go conditions for accuracy and reaction times (Supplemental Table 1). The only exception was for the Fearful No-go conditions for which healthy control group showed greater accuracy than the BDD group. However, to further make sure that accuracy or reaction times were not driving any of the imaging results we used them both as covariates in all second level imaging analysis.

### Condition Effects

The effect of each of the following conditions: Happy face inhibition (Happy Nogo – Neutral Go), Fearful face inhibition (Fearful Nogo-Neutral Go), Emotional face inhibition ((Happy or Fear) Nogo – Neutral Go) and Non-emotional face inhibition ((Male or Female) Nogo – (Female or Male) Go) are presented in Supplementary figure S-2 and Supplementary Table 2. Average effect of condition was examined at voxel-wise threshold of p < .001 (uncorrected) and masks at cluster-wise corrected significance of p < .05 (calculated using 3dClustsim’s auto-correlation function (ACF)) were made at k ≥ 86 voxels (happy face inhibition), 66 voxels (fearful face inhibition), 83 voxels (emotional face inhibition), 95 voxels (non-emotional face inhibition), 102 voxels (happy face inhibition vs. non-emotional face inhibition), 75 voxels (fearful face inhibition vs. non-emotional face inhibition), and 86 voxels (emotional face inhibition vs. non-emotional face inhibition). Average effects of condition masks were used for group analysis as described below.

### Within-Group Imaging Results

We conducted one sample t-tests for each relevant group and visualized differentially activated clusters masked for average effects of the condition in each of the groups for the each of the conditions. AFNI’s 3dClustSim with ACF parameters was used to estimate the cluster size threshold for clusters at voxel-wise threshold of p < .005 (uncorrected) for the MDD+, MDD− and HC groups which would represent the cluster-wise corrected significance of p < 0.05. Due to smaller size of BDD group, voxel-wise threshold was increased to p < .01 for graphical purposes (Supplemental Figures S-3 to S-9). Within-group imaging results are detailed in the Supplemental Material.

Next, we conducted a linear mixed effects univariate analysis to identify between group differences.

### Between-Group Imaging Results

For main effects for group × condition, AFNI’s 3dClustSim with ACF parameters was used to estimate the cluster size threshold for clusters at voxel-wise threshold of p < .001 (uncorrected) which would represent the cluster-wise corrected significance of p < .01. The cluster size threshold was calculated to be k = 21 voxels (happy face inhibition), 60 voxels (fearful face inhibition), 70 voxels (emotional face inhibition), 57 voxels (non-emotional face inhibition), 61 voxels (happy face inhibition vs. non-emotional face inhibition), 42 voxels (fearful face inhibition vs. non-emotional face inhibition), and 39 voxels (emotional face inhibition vs. non-emotional face inhibition). Significant results are depicted in Table 2. Also, presented are post-hoc pairwise group comparisons at p < .05 corrected for multiple comparisons.

#### Happy Face Inhibition

There were no significant main effects for group × condition during happy face inhibition.

#### Fearful Face Inhibition

There were no significant main effects for group × condition during fearful face inhibition.

#### Emotional Face Inhibition

There was significant main effects for group × condition in right cuneus, with post-hoc analysis showing lower activation in the MDD− group compared to the MDD+ and BDD groups (Supplemental Figure S-10(a)).

#### Non-Emotional Face Inhibition

There was significant main effect for group × condition in right temporal pole which had lower activation in the HC group compared to the MDD− but showed no difference with the BDD group and the MDD+ group (Supplemental Figure S-10(b)).

Contrast between emotional conditions vs. the non-emotional condition revealed more extensive differences between groups.

#### Happy Face Inhibition vs. Non-Emotional Face Inhibition

There was significant main effect for group × condition in right angular gyrus, right medial temporal pole, right precuneus, right middle frontal gyrus, right temporal pole, and left inferior temporal gyrus as shown in Supplemental Figure S-11. In these regions, the left inferior temporal gyrus showed lower activation in the MDD− group compared to the MDD+ and HC groups. Moreover, right angular gyrus and the right precuneus showed lower activation in the MDD− group compared to the BDD and HC groups.

#### Fearful Face Inhibition vs. Non-Emotional Face Inhibition

There was significant main effect for group × condition in right medial temporal pole with post-hoc analyses showing lower activation in the MDD− group compared to the BDD and MDD+ groups as shown in Supplemental Figure S-12.

#### Emotional Face Inhibition vs. Non-Emotional Face Inhibition

This comparison showed more extensive differences between the groups. There were significant main effects for group × condition in right posterior cingulate cortex, right precuneus, right medial temporal pole, right angular gyrus, right hippocampus (within 4mm), right superior frontal gyrus, right fusiform gyrus, right superior temporal gyrus, right cerebellum (IX), and left inferior temporal gyrus as shown in Figure 1. Most of these significant regions showed lower activation in the MDD− group compared to the BDD and MDD+ groups.

**Figure 1.**
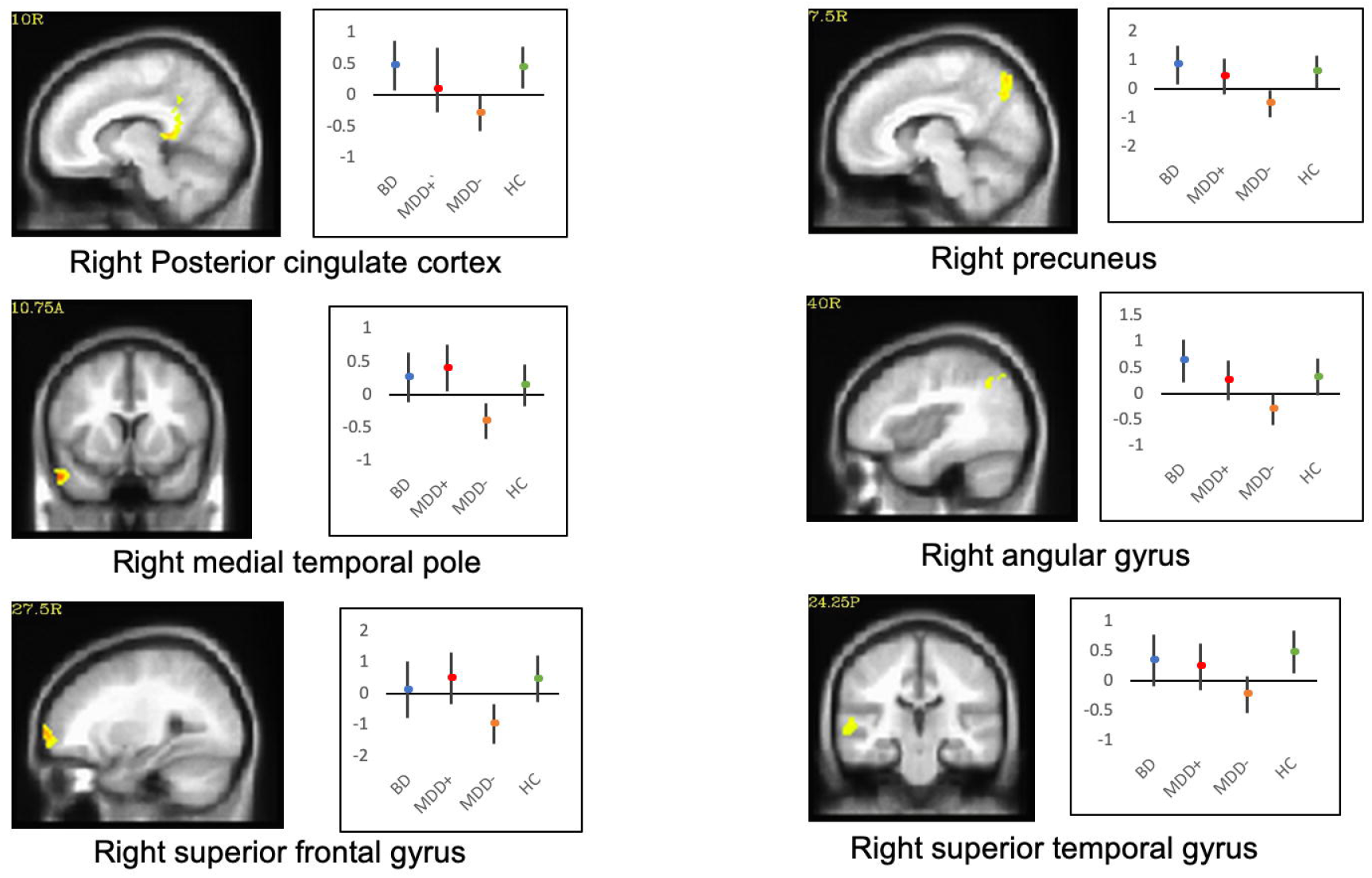
Group activation differences during Emotional face inhibition vs. Non-Emotional face inhibition. The figures are shown at cluster size K ≥ 39 voxels, cluster-wise corrected significance of p < .01. Mean and 95% confidence intervals for average activity within main effect cluster are represented. There were significant main effects for group X condition in 11 ROIs. This figure shows 6 of those ROIs.

Bipolar I vs Bipolar II: as some studies have reported differences in functional imaging parameters between bipolar I and II subtypes we conducted a preliminary analysis of the differences between the two groups in the post-hoc analysis. No corrected significant results were found between the two groups.

Next multivariate machine learning was applied to investigate the individual-level classification accuracy of the Go-No-go task for subjects in each group (Frangou, Dima et al. 2017).

### Classification Results

Classification of beta images using GPC were performed only on the contrast which showed significant main effects for group × condition at cluster-wise corrected significance p < .01 as described above. Classification results are detailed in Table 3. Receiver Operating Characteristics (ROC) curves with Area Under Curve (AUC) of classification are depicted from Figure 2 and Supplementary Figure S-13 and S-14, along with discrimination weight maps.

**Table 3.** Classification Results between BDD, MDD+, and MDD− Based on Beta Images Masked by Average Effect of Condition Mask Using Gaussian Process Classifier.

|  | BDD vs. MDD- |  |  | BDD vs. MDD+ |  |  | MDD+ vs. MDD- |  |  | (BDD & MDD+) vs. MDD- |  |  | BDD vs. (MDD+ & MDD-) |  |  |
| --- | --- | --- | --- | --- | --- | --- | --- | --- | --- | --- | --- | --- | --- | --- | --- |
|  | Happy Face Inhibition vs. Non-Emotional Face Inhibition | Fearful Face Inhibition vs. Non-Emotional Face Inhibition | Emotional Face Inhibition vs. Non-Emotional Face Inhibition | Happy Face Inhibition vs. Non-Emotional Face Inhibition | Fearful Face Inhibition vs. Non-Emotional Face Inhibition | Emotional Face Inhibition vs. Non-Emotional Face Inhibition | Happy Face Inhibition vs. Non-Emotional Face Inhibition | Fearful Face Inhibition vs. Non-Emotional Face Inhibition | Emotional Face Inhibition vs. Non-Emotional Face Inhibition | Happy Face Inhibition vs. Non-Emotional Face Inhibition | Fearful Face Inhibition vs. Non-Emotional Face Inhibition | Emotional Face Inhibition vs. Non-Emotional Face Inhibition | Happy Face Inhibition vs. Non-Emotional Face Inhibition | Fearful Face Inhibition vs. Non-Emotional Face Inhibition | Emotional Face Inhibition vs. Non-Emotional Face Inhibition |
| **AUC** | 0.73 | 0.66 | 0.70 | 0.66 | 0.69 | 0.68 | 0.62 | 0.66 | 0.72 | 0.70 | 0.63 | 0.71 | 0.65 | 0.66 | 0.53 |
| **Accuracy** | 74 | 67 | 75 | 58 | 67 | 63 | 62 | 70 | 70 | 66 | 57 | 65 | 74 | 75 | 69 |
| **Balanced Acc.** | 70 | 56 | 70 | 56 | 66 | 61 | 60 | 65 | 67 | 66 | 58 | 65 | 58 | 59 | 55 |
| **Sensitivity** | 60 | 25 | 55 | 45 | 60 | 55 | 50 | 39 | 54 | 65 | 50 | 65 | 30 | 30 | 30 |
| **Specificity** | 81 | 88 | 85 | 68 | 71 | 68 | 71 | 90 | 80 | 68 | 66 | 66 | 87 | 88 | 80 |
| **PPV** | 60 | 50 | 65 | 50 | 60 | 55 | 54 | 73 | 65 | 70 | 63 | 69 | 40 | 43 | 30 |
| **NPV** | 81 | 71 | 80 | 63 | 71 | 68 | 67 | 69 | 72 | 62 | 53 | 61 | 81 | 81 | 80 |
| **p-value** | 0.005 | 0.018 | 0.006 | 0.237 | 0.033 | 0.091 | 0.078 | 0.026 | 0.015 | 0.005 | 0.114 | 0.012 | 0.076 | 0.041 | 0.172 |

BDD, bipolar depressed group; MDD+, major depressive disorder patients with subthreshold mania symptoms; MDD-, major depressive disorder patients with no subthreshold mania symptoms or risk factors for BD; AUC, area under the receiver operating characteristics (ROC) curve; Balanced Acc.; balanced accuracy; PPV, positive predictive value; NPV, negative predictive value

**Figure 2.**
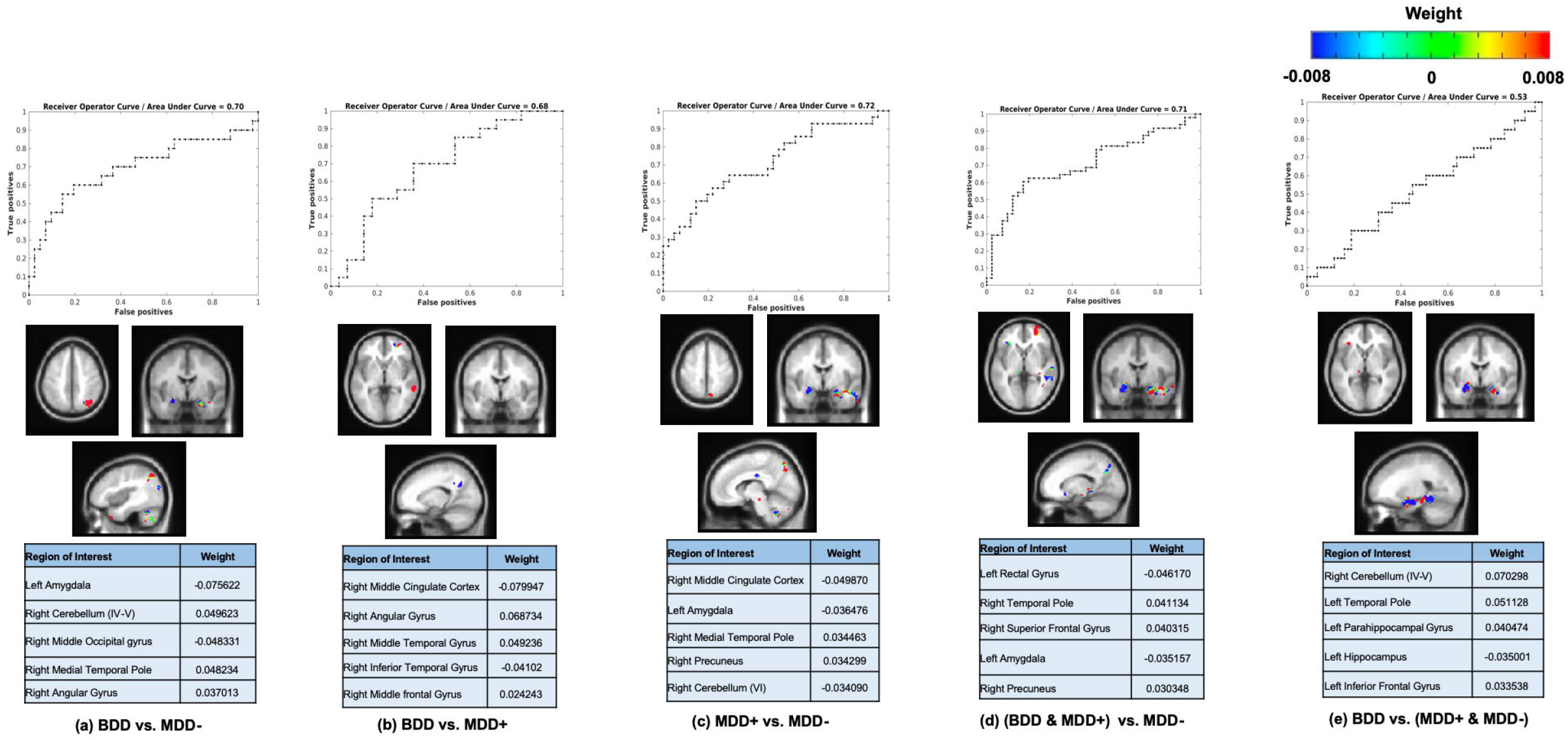
Receiver Operating Characteristics (ROC) Curves with Area Under Curve (AUC) and color weighted discrimination maps of classification for emotional face inhibition vs. non-emotional face inhibition generated by classifier using beta images masked by average effect of condition mask and top five most weighted brain regions for classification. (a) BDD vs. MDD− classification. Positive weights (red) represent the voxels contributing to classification as a BDD subject, while negative weights (blue) represent the voxels contributing to classification as an MDD− subject. (b) MDD+ vs. MDD− classification. Positive class: MDD+ subject, negative class : MDD− subject. (c) (BDD & MDD+) vs. MDD− classification. Positive class : (BDD & MDD+) subject, negative class: MDD− subject. (d) BDD vs.(MDD+ & MDD−) classification. Positive class : BDD subject, negative class : (MDD+ & MDD−) subject.

#### Happy Face Inhibition vs. Non-Emotional Face Inhibition

Classifier using beta images for the happy face inhibition vs. non-emotional inhibition contrast showed significant discrimination (AUC: 0.73; balanced accuracy: 70% (*p* = 0.005)) between BDD vs. MDD− subjects. The largest discriminating clusters were in temporal and parietal regions. The classifier between (BDD & MDD+) and MDD− groups also presented significant discrimination (AUC: 0.70; balanced accuracy: 66% (*p* = 0.005)). Discrimination maps showed that the temporal and parietal brain regions contributed most to these classifiers as shown in Supplemental Figure S-13.

#### Fearful Face Inhibition vs. Non-Emotional Face Inhibition

The fearful face inhibition vs. non-emotional inhibition contrast (AUC: 0.66; balanced accuracy: 65% (*p* = 0.026)) showed significant discrimination between MDD+ and MDD− . The classifier between BDD and (MDD+ & MDD−) groups also provided the significant discrimination (AUC: 0.66; balanced accuracy: 59% (*p* = 0.041)). Discrimination maps showed that the temporal and parietal brain regions contributed most to these classifiers as shown in Supplemental Figure S-14.

#### Emotional Face Inhibition vs. Non-Emotional Face Inhibition

Emotional inhibition vs. non-emotional inhibition contrast (AUC: 0.70; balanced accuracy: 70% *(p* = 0.006)) also showed significant discrimination between BDD and MDD− . The classifier between MDD+ and MDD− groups provided the best discrimination (AUC: 0.72; balanced accuracy: 67% (*p* = 0.015)). The emotional inhibition vs. non-emotional inhibition contrast (AUC: 0.71; balanced accuracy: 65% (*p* = 0.012)) also showed significant discrimination between (BDD & MDD+) versus MDD− . Discrimination maps showed that the temporal and parietal brain regions contributed most to these classifiers. In addition, the left amygdala region contributed to all these classification of the MDD− group for all pairs as shown in Figure 1.

#### Change in classification of MDD group over follow-up

Three MDD subjects (one MDD+ and two MDD−) converted to BD diagnosis over course of follow-up (Koirala, Hu et al. 2019). We examined the original diagnostic label and the predicted diagnosis by classifier explained in the section multivariate pattern classification. One MDD− subject who converted was classified as BDD group in the classifier between BDD and MDD− for the happy face inhibition vs. non-emotional inhibition contrast. Another MDD− subject who converted was classified as the MDD+ group in both the classifiers for happy face inhibition vs. non-emotional inhibition contrast and emotional face inhibition vs. non-emotional inhibition contrast between MDD+ and MDD− . The third subject who converted was an MDD+ subject who was also classified as an MDD+ subject.

## Discussion

In this first study of comparison between medication-free young depressed subjects with and without subthreshold bipolar symptoms and further comparison with diagnosed young bipolar subjects and healthy controls, we found that MDD− group exhibited a different pattern of activation compared to BDD and MDD+ groups. The differences were significant and extensive when emotional response conditions (happy, fearful and combined) were contrasted with non-emotional response condition (gender). Post-hoc analysis of main effect of group revealed that in the MDD− group compared to the BDD and MDD+ groups, exhibited a lower activation in areas of the parietal, temporal and frontal cortices during emotional (happy, fear, and combined emotional) response-inhibition vs non-emotional response-inhibition (Table 2). Importantly, the BDD and MDD+ groups were not significantly different from each other.

The cuneus and adjacent areas of the parietal cortex and the temporal cortex are involved in cognitive appraisal of emotion (Lettieri, Handjaras et al. 2019). Furthermore, the inferior frontal gyrus and cerebellum where lower activations were also seen are associated with motor response-inhibition to emotional stimuli (Wessa, Houenou et al. 2007, Simmonds, Pekar et al. 2008). Response-inhibition to emotional stimuli in MDD− subjects was associated with lowering of activation in the parieto-temporal-frontal emotional appraisal areas. The reason for this widespread deactivation in depression can only be speculated upon. For Go/No-go response-inhibition and other task in which attentional effort is required, studies have reported that either increased or decreased activations of brain areas involved with successful performance on the task may be seen (Lawrence, Ross et al. 2003, Hester, Murphy et al. 2004). The decrease in activation may be necessary to suppress areas of the brain which are overactive due to monitoring of negative internal mood states. The BDD and MDD+ groups were also depressed and would have been expected to show a similar pattern to MDD− subjects. The higher activation in BDD and MDD+ seen in the parieto-temporal-frontal cortex suggests possible increased emotional reactivity, particularly to happy emotional stimuli in BDD and MDD+ groups, as has also been reported in other studies of fMRI activation to emotional stimuli in BD (Blumberg, Donegan et al. 2005, Almeida, Versace et al. 2009, Marchand, Lee et al. 2011, Fournier, Keener et al. 2013).

Importantly, in accordance of the aim and hypothesis of this study we were able to identify a different pattern of activation in BDD and MDD+ group compared to the MDD− group. Therefore, the pattern could potentially be used to accurately classify BDD and MDD− and MDD+ and MDD− subjects. Indeed, machine learning classification of the different patient groups was possible using this difference of activation during emotional inhibition.

Activation related to happy face and combined emotion face response-inhibition vs non-emotional face inhibition showed efficient classification (AUC: .73 and .70) between BDD and MDD− subjects. Combined emotional face response-inhibition vs non-emotional face inhibition also showed good classification properties between MDD+ and MDD− groups (AUC: .70). This classification performance suggests that fMRI activation in response to emotional stimuli could potentially serve as a discriminator for identifying MDD+ subjects. Generally, an AUC of 0.7 to 0.8 is considered suitable classification performance, 0.8 to 0.9 is considered excellent classification performance, and more than 0.9 is considered outstanding classification performance (Mandrekar 2010). Classification results of this study will therefore fall under suitable classification and future studies with more discriminating task design and larger group of subject will need to be conducted.

In combined emotional vs. non-emotional inhibition contrast, notably, the classification contribution of the MDD− group was weighted for amygdala activation pattern (Figure 1). Notably, in the univariate analysis no differences in amygdala activation pattern were found. This finding underscores the importance of multivariate analysis which can reveal findings not seen with univariate analysis. Abnormal amygdala activation has been reported in MDD (Anand, Li et al. 2005) and studies have reported differing activation patterns in MDD and BD (Almeida, Versace et al. 2010, Korgaonkar, Erlinger et al. 2019).

Notably, the post-hoc results of main effects of group was not able to differentiate between BDD and MDD+ groups but there were several instances of significant differences between MDD+ and MDD− groups (Table 2). These findings indicated that the MDD+ group are more similar to the BDD group than the MDD− group. Further support was found due to a stronger discrimination between the combined BDD and MDD+ group compared to the MDD− group (AUC: .70 and .71 for happy and combined emotion vs non-emotional response-inhibition) compared to the BDD group and the combined MDD+ and MDD− groups (traditionally termed unipolar depression) (all AUCs <.70). These findings underscore the importance of also (de Almeida and Phillips 2013) studying MDD+ and MDD− groups separately when comparing bipolar and unipolar depression.

There are several limitations of the current study. While the medication-free status of the participants in this study is a rare strength, it also poses possible selection bias in that the participant population may over represent subjects with more mild conditions, and under represent those where the severity of the disorder is such that medication-free study inclusion would be unethical. Because medication-free psychiatric neuroimaging is greatly understudied, these data are valuable, but may lack full generalizability to medicated patient populations. These results must be compared with future studies in medicated and unmedicated participants to be appropriately generalized. Variances were equal between groups, but the smaller sample size in the bipolar group may have led to lower power for post hoc tests, possibly resulting in more missed conclusions involving the BDD group. For the same reason, we were unable to separate the BDD group into bipolar I and II subtypes for similarities and differences with the MDD+ group. However, we did not find any significant differences between BDI and BDII when the BD group was split in the LME analysis. However, a larger number of subjects in each group need to be studied. In future studies with a larger sample of bipolar subtypes such an investigation will need to be undertaken.

For the classification analyses, ideally a totally separate test-dataset from the training data set should be used, though in this case we did do the leave-one-out per group cross validation. For each iteration of cross validation, one subject from each group was excluded and then was allocated to the test set. In future studies, the classifier developed in this study can be tested using an independent sample. Finally, the classifiers obtained in these analyses could be used to predict the development of BD over time in MDD subjects. In this study, among three MDD subjects converted to BD over time, two of the converted MDD− subject was classified as BDD and the other converted MDD+ subject was classified as MDD+. These results though small provide some face validity to the classifiers. Future studies with a larger number of subjects and longer period of times will need to be conducted to further validate the emotional vs. non-emotional Go-Nogo task as a predict of future conversion to BD in young MDD subjects.

## Conclusion

The findings of the study indicate that the MDD− group was different from both BDD as well as MDD+ young adult groups on the regional activation pattern of emotional vs. non-emotional response inhibition tasks. MDD+ and BDD groups were similar to each other. Future studies need to be conducted to further refine this classifier to predict development of bipolar disorder in young adults presenting with depression.

## FUNDING AND DISCLOSURES

This project was funded by the NIMH to AA (R01MH093420). Dr. Anand, Dr. Altinay, Dr. Koirala, Sidra Speaker, Jungwon Cha and Harish Karne each reported no financial interests or potential conflicts of interest.

## Notes

### Competing Interest Statement

The authors have declared no competing interest.

## REFERENCES

Almeida, J. R. C., A. Versace, S. Hassel, D. J. Kupfer and M. L. Phillips (2010). “Elevated Amygdala Activity to Sad Facial Expressions: A State Marker of Bipolar but Not Unipolar Depression.” Biological Psychiatry 67(5): 414–421.

Almeida, J. R. C. d., A. Versace, A. Mechelli, S. Hassel, K. Quevedo, D. J. Kupfer and M. L. Phillips (2009). “Abnormal Amygdala-Prefrontal Effective Connectivity to Happy Faces Differentiates Bipolar from Major Depression.” Biological Psychiatry 66(5): 451–459.

Amit Etkin, M.D., Ph.D. (2019). “A Reckoning and Research Agenda for Neuroimaging in Psychiatry.” American Journal of Psychiatry 176(7): 507–511.

Anand, A., Y. Li, Y. Wang, J. Wu, S. Gao, L. Bukhari, V. Mathews, A. Kalnin and M. Lowe (2005). “Activity and connectivity of brain mood regulating circuit in depression: A functional magnetic resonance study.” Biological Psychiatry 57(10): 1079–1088.

Angst, J., A. Gamma, F. Benazzi, V. Ajdacic, D. Eich and W. Rössler (2003). “Toward a redefinition of subthreshold bipolarity: epidemiology and proposed criteria for bipolar-Ĳ, minor bipolar disorders and hypomania.” Journal of Affective Disorders 73(1): 133–146.

Blumberg, H. P., N. H. Donegan, C. A. Sanislow, S. Collins, C. Lacadie, P. Skudlarski, R. Gueorguieva, R. K. Fulbright, T. H. McGlashan, J. C. Gore and J. H. Krystal (2005). “Preliminary evidence for medication effects on functional abnormalities in the amygdala and anterior cingulate in bipolar disorder.” Psychopharmacology (Berl) 183(3): 308–313.

Brodersen, K. H., F. Haiss, C. S. Ong, F. Jung, M. Tittgemeyer, J. M. Buhmann, B. Weber and K. E. Stephan (2011). “Model-based feature construction for multivariate decoding.” Neuroimage 56(2): 601–615.

Chen, G., Z. S. Saad, J. C. Britton, D. S. Pine and R. W. Cox (2013). “Linear mixed-effects modeling approach to FMRI group analysis.” Neuroimage 73: 176–190.

Chen, G., P. A. Taylor and R. W. Cox (2017). “Is the statistic value all we should care about in neuroimaging?” NeuroImage 147: 952–959.

Coryell, W., J. Endicott, J. D. Maser, M. B. Keller, A. C. Leon and H. S. Akiskal (1995). “Longterm stability of polarity distinctions in the affective disorders.” The American journal of psychiatry 152(3): 385–390.

Cox, R. W., G. Chen, D. R. Glen, R. C. Reynolds and P. A. Taylor (2017). “FMRI Clustering in AFNI: False-Positive Rates Redux.” Brain Connect 7(3): 152–171.

de Almeida, J. R. C. and M. L. Phillips (2013). “Distinguishing between Unipolar Depression and Bipolar Depression: Current and Future Clinical and Neuroimaging Perspectives.” Biological Psychiatry 73(2): 111–118.

Fiedorowicz, J. G., J. Endicott, A. C. Leon, D. A. Solomon, M. B. Keller and W. H. Coryell (2011). “Subthreshold hypomanic symptoms in progression from unipolar major depression to bipolar disorder.” The American journal of psychiatry 168(1): 40–48.

Forman, S. D., J. D. Cohen, M. Fitzgerald, W. F. Eddy, M. A. Mintun and D. C. Noll (1995). “Improved assessment of significant activation in functional magnetic resonance imaging (fMRI): use of a cluster-size threshold.” Magn Reson Med 33(5): 636–647.

Fournier, J. C., M. T. Keener, B. C. Mullin, D. M. Hafeman, E. J. Labarbara, R. S. Stiffler, J. Almeida, D. M. Kronhaus, E. Frank and M. L. Phillips (2013). “Heterogeneity of amygdala response in major depressive disorder: the impact of lifetime subthreshold mania.” Psychol Med 43(2): 293–302.

Frangou, S., D. Dima and J. Jogia (2017). “Towards person-centered neuroimaging markers for resilience and vulnerability in Bipolar Disorder.” Neuroimage 145(Pt B): 230–237.

Golland, P. and B. Fischl (2003). “Permutation tests for classification: towards statistical significance in image-based studies.” Inf Process Med Imaging 18: 330–341.

Han, K. M., D. De Berardis, M. Fornaro and Y. K. Kim (2019). “Differentiating between bipolar and unipolar depression in functional and structural MRI studies.” Prog Neuropsychopharmacol Biol Psychiatry 91: 20–27.

Hester, R. L., K. Murphy, J. J. Foxe, D. M. Foxe, D. C. Javitt and H. Garavan (2004). “Predicting success: patterns of cortical activation and deactivation prior to response inhibition.” J Cogn Neurosci 16(5): 776–785.

Hummer, T. A., L. A. Hulvershorn, H. S. Karne, A. D. Gunn, Y. Wang and A. Anand (2013). “Emotional response inhibition in bipolar disorder: a functional magnetic resonance imaging study of trait-and state-related abnormalities.” Biol Psychiatry 73(2): 136–143.

Kling, L. R., K. L. Bessette, S. R. DelDonno, K. A. Ryan, W. C. Drevets, M. G. McInnis, M. L. Phillips and S. A. Langenecker (2018). “Cluster analysis with MOODS-SR illustrates a potential bipolar disorder risk phenotype in young adults with remitted major depressive disorder.” Bipolar Disord 20(8): 697–707.

Koirala, P., B. Hu, M. Altinay, M. Li, A. L. DiVita, K. A. Bryant, H. S. Karne, J. G. Fiedorowicz and A. Anand (2019). “Sub-threshold bipolar disorder in medication-free young subjects with major depression: Clinical characteristics and antidepressant treatment response.” J Psychiatr Res 110: 1–8.

Korgaonkar, M. S., M. Erlinger, I. A. Breukelaar, P. Boyce, P. Hazell, C. Antees, S. Foster, S. M. Grieve, L. Gomes, L. M. Williams, A. W. F. Harris and G. S. Malhi (2019). “Amygdala Activation and Connectivity to Emotional Processing Distinguishes Asymptomatic Patients With Bipolar Disorders and Unipolar Depression.” Biological Psychiatry: Cognitive Neuroscience and Neuroimaging 4(4): 361–370.

Lawrence, N. S., T. J. Ross, R. Hoffmann, H. Garavan and E. A. Stein (2003). “Multiple neuronal networks mediate sustained attention.” J Cogn Neurosci 15(7): 1028–1038.

Lettieri, G., G. Handjaras, E. Ricciardi, A. Leo, P. Papale, M. Betta, P. Pietrini and L. Cecchetti (2019). “Emotionotopy in the human right temporo-parietal cortex.” Nature Communications 10(1): 5568.

Mandrekar, J. N. (2010). “Receiver Operating Characteristic Curve in Diagnostic Test Assessment.” Journal of Thoracic Oncology 5(9): 1315–1316.

Marchand, W. R., J. N. Lee, C. Garn, J. Thatcher, P. Gale, S. Kreitschitz, S. Johnson and N. Wood (2011). “Aberrant emotional processing in posterior cortical midline structures in bipolar II depression.” Prog Neuropsychopharmacol Biol Psychiatry 35(7): 1729–1737.

Merikangas, K. R., R. Jin, J. P. He, R. C. Kessler, S. Lee, N. A. Sampson, M. C. Viana, L. H. Andrade, C. Hu, E. G. Karam, M. Ladea, M. E. Medina-Mora, Y. Ono, J. Posada-Villa, R. Sagar, J. E. Wells and Z. Zarkov (2011). “Prevalence and correlates of bipolar spectrum disorder in the world mental health survey initiative.” Arch Gen Psychiatry 68(3): 241–251.

Nichols, T. E. and A. P. Holmes (2002). “Nonparametric permutation tests for functional neuroimaging: a primer with examples.” Hum Brain Mapp 15(1): 1–25.

Noirhomme, Q., D. Lesenfants, F. Gomez, A. Soddu, J. Schrouff, G. Garraux, A. Luxen, C. Phillips and S. Laureys (2014). “Biased binomial assessment of cross-validated estimation of classification accuracies illustrated in diagnosis predictions.” Neuroimage Clin 4: 687–694.

Nurnberger, J. I., Jr., M. McInnis, W. Reich, E. Kastelic, H. C. Wilcox, A. Glowinski, P. Mitchell, C. Fisher, M. Erpe, E. S. Gershon, W. Berrettini, G. Laite, R. Schweitzer, K. Rhoadarmer, V. V. Coleman, X. Cai, F. Azzouz, H. Liu, M. Kamali, C. Brucksch and P. O. Monahan (2011). “A high-risk study of bipolar disorder. Childhood clinical phenotypes as precursors of major mood disorders.” Arch Gen Psychiatry 68(10): 1012–1020.

Phillips, M. L. and H. A. Swartz (2014). “A critical appraisal of neuroimaging studies of bipolar disorder: toward a new conceptualization of underlying neural circuitry and a road map for future research.” Am J Psychiatry 171(8): 829–843.

Piguet, C., L. Fodoulian, J. M. Aubry, P. Vuilleumier and J. Houenou (2015). “Bipolar disorder: Functional neuroimaging markers in relatives.” Neurosci Biobehav Rev 57: 284–296.

Rasmussen, C. E. (2003). Gaussian processes in machine learning. Summer School on Machine Learning, Springer.

Redlich, R., J. J. Almeida, D. Grotegerd, N. Opel, H. Kugel, W. Heindel, V. Arolt, M. L. Phillips and U. Dannlowski (2014). “Brain morphometric biomarkers distinguishing unipolar and bipolar depression. A voxel-based morphometry-pattern classification approach.” JAMA Psychiatry 71(11): 1222–1230.

Redlich, R., K. Dohm, D. Grotegerd, N. Opel, P. Zwitserlood, W. Heindel, V. Arolt, H. Kugel and U. Dannlowski (2015). “Reward Processing in Unipolar and Bipolar Depression: A Functional MRI Study.” Neuropsychopharmacology 40(11): 2623–2631.

Simmonds, D. J., J. J. Pekar and S. H. Mostofsky (2008). “Meta-analysis of Go/No-go tasks demonstrating that fMRI activation associated with response inhibition is task-dependent.” Neuropsychologia 46(1): 224–232.

Strakowski, S. M., C. M. Adler, J. Almeida, L. L. Altshuler, H. P. Blumberg, K. D. Chang, M. P. DelBello, S. Frangou, A. McIntosh, M. L. Phillips, J. E. Sussman and J. D. Townsend (2012). “The functional neuroanatomy of bipolar disorder: a consensus model.” Bipolar Disord 14(4): 313–325.

Townsend, J. D., S. Y. Bookheimer, L. C. Foland-Ross, T. D. Moody, N. I. Eisenberger, J. S. Fischer, M. S. Cohen, C. A. Sugar and L. L. Altshuler (2012). “Deficits in inferior frontal cortex activation in euthymic bipolar disorder patients during a response inhibition task.” Bipolar Disord 14(4): 442–450.

Vierck, E., R. J. Porter and P. R. Joyce (2015). “Facial recognition deficits as a potential endophenotype in bipolar disorder.” Psychiatry Res 230(1): 102–107.

Wessa, M., J. Houenou, M. L. Paillere-Martinot, S. Berthoz, E. Artiges, M. Leboyer and J. L. Martinot (2007). “Fronto-striatal overactivation in euthymic bipolar patients during an emotional go/nogo task.” Am J Psychiatry 164(4): 638–646.

Zimmermann, P., T. Brückl, A. Nocon, H. Pfister, R. Lieb, H.-U. Wittchen, F. Holsboer and J. Angst (2009). “Heterogeneity of DSM-IV major depressive disorder as a consequence of subthreshold bipolarity.” Archives of general psychiatry 66(12): 1341–1352.

