## Supplemental Materials for "Neuroimaging Correlates of Emotional Response-Inhibition Discriminate Between Young Depressed Adults With and Without Sub-threshold Bipolar Symptoms"

### Inclusion and Exclusion Criteria

All subjects took part in the study after signing an informed consent document approved by the Institutional Review Board at the Cleveland Clinic. Subjects under the age of 18 signed an assent form and a parent signed the consent form. Each participant was paid $75 for screening and $75 for a magnetic resonance imaging scans (MRI). All participants underwent a clinical interview with a psychiatrist, a detailed structured diagnostic interview (Mini Neuropsychiatric Interview (MINI)) and administration of clinical scales for depression and mania.

Depressed participants satisfied DSM-IV-R criteria for BD or MDD in a current depressive episode, with Hamilton Depression Rating Scale (Hamilton 1960) (HDRS) ≥ 15 but ≤ 25 at the time of screening.. Exclusion criteria for all participants included Young Mania Rating Scale (Young, Biggs et al. 1978) (YMRS) > 10 at the time of screening; a lifetime diagnosis of schizophrenia or schizoaffective disorder; a current primary anxiety disorder; psychotropic medication use within the past 2 weeks; fluoxetine use within the past 4 weeks; acute suicidal or homicidal ideation or behavior; recent (<1 week) use of alcohol; current pregnancy or breastfeeding; positive urine toxicology test at baseline; and contraindications to MRI. Additional inclusion criteria for MDD subjects included having never met criteria for mania or hypomania.

Healthy subjects had no personal or family history of psychiatric illness or alcohol or substance abuse/dependence; no current use of any centrally acting medications; no alcohol use in the past week; and no serious medical or neurological illness.

### Imaging Procedure

Participants learned the task prior to the scanning session. The stimulus presentation was programmed in E-Prime 2.0 (Psychology Software Tools, Pittsburg PA), which also recorded reaction time and participant responses by reading inputs from a button box.

After a short scout imaging scan to center the field of view, one high-resolution T1-wieghted 3D magnetization prepared rapid gradient echo (MPRAGE) (1.0 x 1.0 x 1.2 $mm^3$ voxels) was acquired for later use in co-registration and normalization. Subsequent functional scans acquired blood oxygen level dependent (BOLD) signal using a T2*-weighted gradient-echo echo-planar imaging sequence (164 volumes, repetition/echo time 2800/29 msec, 39 slices, 2.5 x 2.5 x 3.5 $mm^3$ voxels). The imaging data was acquired at Cleveland Clinic Main Campus imaging center using a Siemens 3T Trio MR Scanner (Siemens AG, Berlin, Germany) with a 32 receive channel head coil and electronically transferred to the Cleveland Clinic imaging archive system. During the data collection in Cleveland Clinic the MR scanner underwent upgrade from Trio to Prisma. The first 4 volumes of the functional run were discarded due to the T1 stabilization process, and analysis included 160 volumes taken over 7 minutes and 28 seconds.

Functional Neuroimaging Data Analysis: Images were preprocessed and analyzed in AFNI (https://afni.nimh.nih.gov, Cox 1996) according to Analysis of Functional Neuro Images. Subjects were excluded from imaging analysis if translational or rotational movement exceeded 2 mm or 2 in any direction. Functional scans were realigned then co-registered to the anatomical scan and normalized into MNI space. During normalization, images were resampled every 2.5 mm and were smoothed with an 8x8x8 mm Gaussian kernel.

### Permutation test

Permutation testing was performed for each classifier with 1000 permutations, which provided a p-value for the balanced accuracy of classification. A permutation test is a non-parametric test estimating the distribution of the null hypothesis from the data. In particular, for each permutation, assume that there is no class label information. The data was randomly permuted (with respect to the labels) and the balanced accuracy was recomputed with the new labels. Since the permuted labels were random, the new balanced accuracy was expected to reflect the chance distribution. Thus, the p-value was calculated by the sum of all new balanced accuracy which were equal or larger than the balanced accuracy observed when the original labels were used divided by the number of permutations. (Nichols and Holmes 2002, Golland and Fischl 2003, Noirhomme, Lesenfants et al. 2014).

### Within-Group Imaging Results

For the happy face inhibition, the MDD- group showed lower activation within the frontal gyrus, marked lower activation of the orbital gyrus, and lower activation in the anterior cingulate cortex, whereas the MDD+ group and BDD groups had lower activation only in the lateral orbital gyrus.

For the fearful face inhibition, MDD- showed lower activation in lateral orbital and temporal areas, whereas MDD+ and BDD showed only lower activation in the lateral orbital areas.

For the emotional face inhibition (happy and fearful combined), MDD- showed widespread lower activation in frontal, temporal, and parietal cortices, whereas MDD+ and BDD groups showed lower activation mainly in the frontal orbital areas.

Conversely, for the non-emotional face inhibition (gender), MDD+ and BDD groups showed wide-spread lower activation in the orbital, temporal, and parietal areas, whereas MDD- showed increased activation in parietal areas. HC groups had low magnitude activations or deactivations during the different inhibition conditions.

For the happy face inhibition vs non-emotional face inhibition, MDD+ and BDD groups showed increased activation in the frontal, parietal, and right temporal areas, whereas MDD- showed lower activation in parietal and both side of temporal areas.

For the fearful face inhibition vs non-emotional face inhibition, BDD, MDD+ and HC groups showed increased activation in the temporal and parietal areas.

For the emotional face inhibition vs non-emotional face inhibition, BDD, MDD+ and HC groups showed increased activation in the frontal, parietal, and right temporal areas, whereas MDD- showed lower activation in parietal and both side of temporal areas. Moreover, MDD+ showed lower activation in left temporal area.

**Supplementary Table 1.** Task Performance Measures.

| | BDD (%) | MDD+ (%) | MDD- (%) | HC (%) | ANOVA -tests F-value (p-value) | post-hoc test Bonferroni corrected (p < .05) |
|---|---|---|---|---|---|---|
| ***Accuracy Go blocks*** | | | | | | |
| Neutral Go | 83 | 82 | 83 | 84 | 0.168 (0.918) | |
| Male Go | 98 | 96 | 99 | 100 | 1.062 (0.368) | |
| Female Go | 87 | 88 | 85 | 92 | 0.938 (0.425) | |
| ***Accuracy No-Go from the No-Go/Go blocks*** | | | | | | |
| Happy No-go | 94 | 96 | 98 | 99 | 2.744 (0.046) | |
| Fear No-go | 90 | 92 | 95 | 97 | 4.812 (0.003) | BDD < HC |
| Male No-go | 96 | 94 | 96 | 97 | 0.934 (0.427) | |
| Female No-go | 87 | 86 | 87 | 93 | 1.566 (0.202) | |
| ***Reaction Times (msec) Go blocks*** | | | | | | |
| Neutral Go | 548 | 525 | 533 | 567 | 1.140 (0.336) | |
| Male Go | 476 | 466 | 511 | 521 | 2.285 (0.083) | |
| Female Go | 516 | 516 | 506 | 549 | 0.945 (0.421) | |
| ***Reaction Times (msec) Go from the No-Go/Go blocks*** | | | | | | |
| Happy No-go go block | 535 | 506 | 535 | 529 | 0.869 (0.460) | |
| Fear No-go go block | 542 | 512 | 549 | 545 | 1.840 (0.144) | |
| Male No-go go block | 510 | 503 | 520 | 542 | 1.487 (0.222) | |
| Female No-go go block | 530 | 532 | 549 | 557 | 1.307 (0.276) | |

*Degrees of freedom for all tests were (3,114). *ANOVA* tests were performed and post-hoc tests were performed with Bonferroni for multiple comparisons. Reaction times for the Go part of the No-go/Go blocks were used.

- BDD: bipolar depressed group
- MDD+: major depressive disorder patients who are at high risk for developing BD
- MDD- : major depressive disorder patients who are at low risk for developing BD
- HC: Healthy Control

**Supplementary Table 2.** Significant Results of Condition Effects.

| Condition Effects | | | |
|---|---|---|---|
| **Activation Area** | **Cluster Size** | **Peak MNI (X Y Z)** | **Peak F with Degree of Freedom (3, 110)** |
| ***Happy Face Inhibition*** | | | |
| Right Anterior Cingulate Cortex | 256 | 10, 38, 26 | 11.685 |
| Right Superior Frontal Gyrus | 203 | 28, 53, 38 | 8.886 |
| Right Superior Orbital Gyrus | 141 | 20, 51, -14 | 10.124 |
| Right Middle Frontal Gyrus | 114 | 28, 61, 23 | 11.844 |
| ***Fear Face Inhibition*** | | | |
| Left Fusiform Gyrus | 298 | -25, -44, -12 | 10.950 |
| Right Superior Frontal Gyrus | 263 | 22, 68, 16 | 13.169 |
| Right Middle Orbital Gyrus | 191 | 28, 41, -17 | 10.338 |
| Left Precuneus | 128 | -2, -57, 68 | 10.012 |
| Right Cuneus | 125 | 5, -89, 33 | 7.802 |
| Right Fusiform Gyrus | 111 | 22, -47, -12 | 7.539 |
| Right Anterior Cingulate Cortex | 106 | 10, 38, 26 | 8.138 |
| ***Emotional Face Inhibition*** | | | |
| Right Superior Frontal Gyrus | 5933 | 25, 63, 20 | 19.832 |
| Left Fusiform Gyrus | 2592 | -28, -42, -14 | 13.959 |
| Left Precuneus | 1232 | -2, -57, 68 | 13.214 |
| Right Angular Gyrus | 327 | 58, -57, 33 | 9.392 |
| Right Inferior Temporal Gyrus | 287 | 55, -42, -17 | 11.360 |
| Right Cerebellum (Crus 2) | 196 | 8, -87, -37 | 9.749 |
| Left Amygdala | 175 | -30, 1, -24 | 12.660 |
| Left Cerebellum (Crus 2) | 149 | -18, -74, -40 | 8.336 |
| Right Superior Temporal Gyrus | 131 | 58, -17, 3 | 10.113 |
| Left Middle Cingulate Cortex (within 4mm) | 104 | -8, -22, 28 | 14.498 |
| ***Non-Emotional Face Inhibition*** | | | |
| Right Angular Gyrus | 19449 | 50, -52, 33 | 33.333 |
| Left Middle Frontal Gyrus | 673 | -40, 28, 43 | 12.973 |
| Right Inferior Temporal Gyrus | 328 | 42, -62, -7 | 10.909 |
| Left Middle Frontal Gyrus | 224 | -35, 61, 3 | 9.786 |

| | | | |
|---|---|---|---|
| ***Happy Face Inhibition vs. Non-Emotional Face Inhibition*** | | | |
| Right SupraMarginal Gyrus | 4038 | 58, -44, 36 | 19.513 |
| Right Precuneus | 2812 | 12, -47, 38 | 24.003 |
| Right Inferior Frontal Gyrus | 2116 | 50, 46, -17 | 16.075 |
| Left Inferior Temporal Gyrus | 1325 | -62, -49, -17 | 13.866 |
| Right Calcarine Gyrus | 387 | 8, -89, 6 | 11.310 |
| Left Amygdala | 129 | -22, 1, -22 | 11.250 |
| Left Superior Frontal Gyrus | 108 | -20, 16, 63 | 10.874 |
| ***Fear Face Inhibition vs. Non-Emotional Face Inhibition*** | | | |
| Right SupraMarginal Gyrus | 3115 | 58, -44, 36 | 19.369 |
| Right Inferior Frontal Gyrus | 1785 | 50, 46, -17 | 14.206 |
| Right Precuneus | 1402 | 12, -47, 38 | 18.535 |
| Left Inferior Temporal Gyrus | 690 | -62, -49, -17 | 12.828 |
| Left Inferior Parietal Lobule | 371 | --55, -44, 53 | 13.909 |
| Right Calcarine Gyrus | 295 | 10, -92, 3 | 12.429 |
| Left Cerebellum (Crus 2) | 275 | -45, -62, -47 | 10.658 |
| Right Medial Temporal Pole | 149 | 58, 8, -30 | 12.119 |
| ***Emotional Face Inhibition vs. Non-Emotional Face Inhibition*** | | | |
| Right Medial Temporal Pole | 2378 | 58, 11, -30 | 13.407 |
| Right Precuneus | 1924 | 10, -49, 38 | 16.055 |
| Left Inferior Temporal Gyrus | 394 | -62, -49, -17 | 11.551 |
| Left Amygdala | 311 | -25, 3, -22 | 12.122 |
| Cerebellar Vermis (9) | 215 | 2, -59, -37 | 10.993 |
| Right Hippocampus | 197 | 22, -2, -22 | 11.581 |
| Right Superior Frontal Gyrus | 185 | 30, 66, 0 | 11.177 |
| Left Parahippocampal Gyrus | 160 | -22, -34, -12 | 9.273 |

**Supplementary Table 3.** Significant Results of Main Effects for Group x Condition of the three groups and Post-hoc Comparisons with Bonferroni Correction for Multiple Comparisons.

| **Main Effects for Group x Condition** | | | | **Post-hoc Comparisons** |
|---|---|---|---|---|
| **Activation Area** | **Cluster Size** | **Peak MNI (X Y Z)** | **Peak F with Degree of Freedom (2, 81)** | **Post-hoc test Bonferroni Corrected p < 0.05** |
| ***Emotional Face Inhibition*** | | | | |
| Right Posterior Cingulate Cortex | 147 | 10, -42, 10 | 8.559 | BDD > MDD- |
| Right Cerebellum (Crus 1) | 128 | 45, -57, -27 | 8.637 | BDD > MDD-<br>MDD+ > MDD- |
| Right Cuneus | 70 | 20, -79, 43 | 7.338 | BDD > MDD-<br>MDD+ > MDD- |
| ***Non-Emotional Face Inhibition*** | | | | |
| Right Medial Temporal Pole | 41 | 50, 18, -34 | 8.326 | MDD- > BD<br>MDD- > MDD+ |
| Left Inferior Temporal Gyrus | 26 | -65, -49, -17 | 8.805 | MDD- > MDD+ |
| ***Happy Face Inhibition vs. Non-Emotional Face Inhibition*** | | | | |
| Right Medial Temporal Pole | 256 | 58, 11, -30 | 12.933 | MDD+ > MDD- |
| Right Precuneus | 135 | 10, -74, 56 | 8.120 | BDD > MDD-<br>MDD+ > MDD- |
| Right Angular Gyrus | 95 | 45, -69, 40 | 7.030 | BDD > MDD-<br>MDD+ > MDD- |
| Right Fusiform Gyrus | 51 | 42, -27, -30 | 7.629 | MDD+ > MDD- |
| Right Inferior Temporal Gyrus | 50 | 60, -42, -17 | 7.514 | MDD+ > MDD- |
| Right Hippocampus (within 4mm) | 30 | 40, -4, -20 | 7.761 | MDD+ > MDD- |
| Right Superior Parietal Lobule | 20 | 32, -69, 56 | 6.358 | BDD > MDD-<br>MDD+ > MDD- |
| Right Superior Frontal Gyrus | 17 | 28, 68, 3 | 7.582 | none |
| ***Fear Face Inhibition vs. Non-Emotional Face Inhibition*** | | | | |
| Right Medial Temporal Pole | 61 | 58, 8, -30 | 8.359 | MDD+ > MDD- |
| Right Fusiform Gyrus | 43 | 28, 18, -47 | 7.400 | MDD+ > MDD- |
| ***Emotional Face Inhibition vs. Non-Emotional Face Inhibition*** | | | | |
| Right Medial Temporal Pole | 324 | 58, 11, -30 | 11.911 | BDD > MDD-<br>MDD+ > MDD- |
| Right Posterior Cingulate Cortex | 186 | 10, -37, 8 | 8.624 | BDD > MDD- |
| Right Precuneus | 166 | 8, -72, 53 | 9.007 | BDD > MDD- |
| Right Angular Gyrus | 123 | 40, -57, 38 | 6.967 | BDD > MDD- |
| Right Fusiform Gyrus | 77 | 28, 16, -44 | 9.797 | MDD+ > MDD- |
| Right Cerebellum (IX) | 51 | 8, -57, -47 | 8.120 | BDD > MDD-<br>BDD > MDD+ |
| Right Superior Occipital Gyrus | 41 | 32, -79, 43 | 6.809 | BDD > MDD- |
| Left Precuneus | 37 | -10, -69, 50 | 7.649 | BDD > MDD- |
| Right Inferior Temporal Gyrus | 30 | 55, -42, -20 | 6.954 | MDD+ > MDD- |
| Right Superior Frontal Gyrus | 29 | 28, 68, 3 | 8.378 | none |
| Right Middle Temporal Gyrus | 18 | 58, -52, 10 | 6.825 | BDD > MDD- |
| Right Precuneus | 16 | 22, -59, 28 | 6.386 | none |

**Supplementary Table 4.** Significant Results of Main Effects for Group x Condition between depressed patients group (BDD & MDD+ & MDD-) and HC using the average effect of condition mask.

| Condition | Activation Area | MNI peak | Peak F with Degree of Freedom (1, 113) | Cluster Size | PA vs. HC |
|---|---|---|---|---|---|
| ***Non-Emotional Face Inhibition*** | Right Temporal Pole | 32, 21, -27 | 7.42 | 114 | PA > HC |
| | Left Cerebellum (VI) | -8, -64, -7 | 6.75 | 72 | PA > HC |
| | Left Cerebellum (Crus 1) | -40, -67, -27 | 6.16 | 48 | PA > HC |
| ***Happy Face Inhibition vs. Non-Emotional Face Inhibition*** | Left Inferior Frontal Gyrus | -38, 16, 20 | 7.02 | 47 | PA < HC |
| | Right Temporal Pole | 30, 16, -27 | 6.37 | 38 | PA < HC |
| | Left Cerebellum (Crus 1) | -40, -71, -22 | 6.33 | 36 | PA < HC |
| ***Emotional Face Inhibition vs. Non-Emotional Face Inhibition*** | Left Temporal Pole | -45, 6, -22 | 6.05 | 16 | PA < HC |
| | Right Temporal Pole | 28, 11, -24 | 5.49 | 11 | PA < HC |
