## Supplemental Figures for "Neuroimaging Correlates of Emotional Response-Inhibition Discriminate Between Young Depressed Adults With and Without Sub-threshold Bipolar Symptoms"

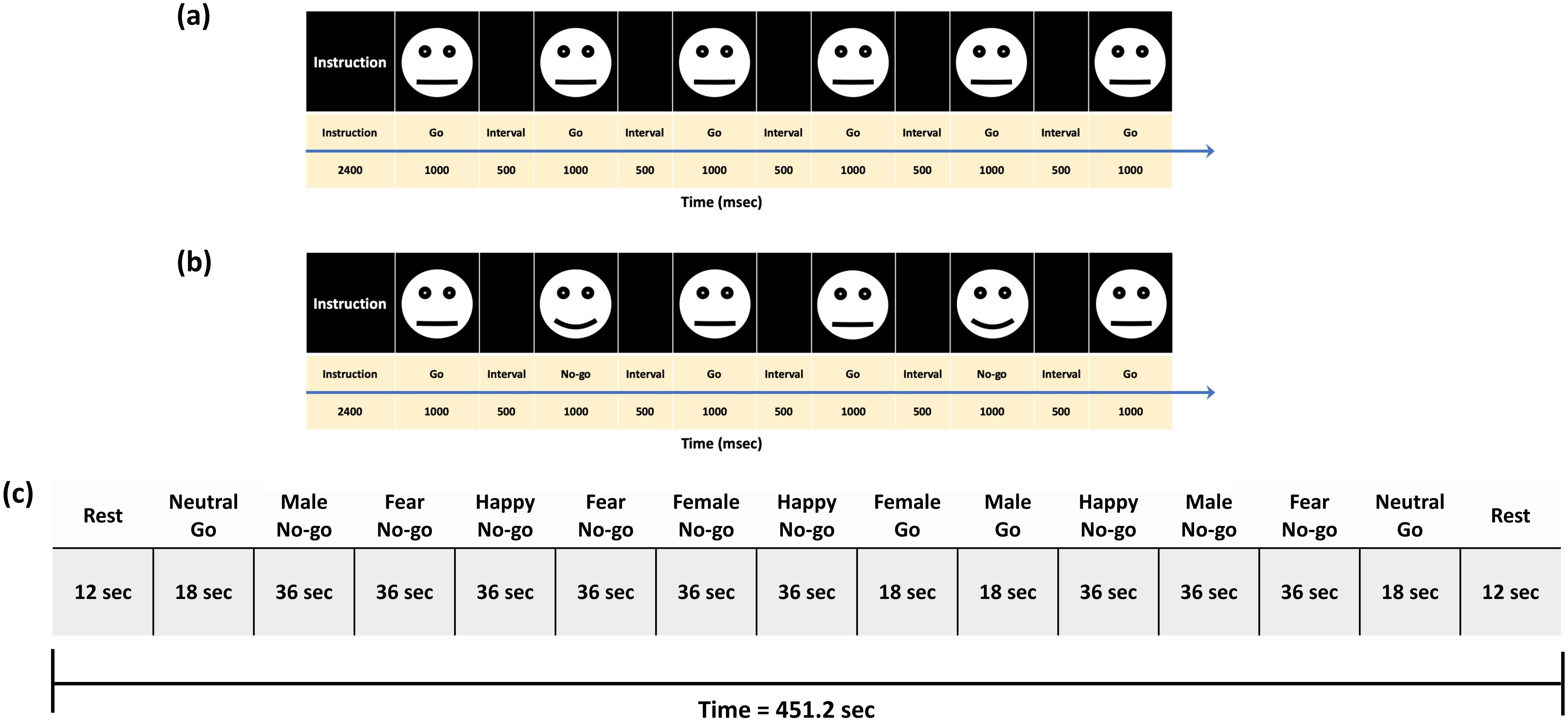


**Figure S-1.** Functional magnetic resonance imaging paradigm. Example of presentation instruction and stimuli for (a) (neutral expression) go and (b) happy no-go blocks. (c) The structure for the entire run.

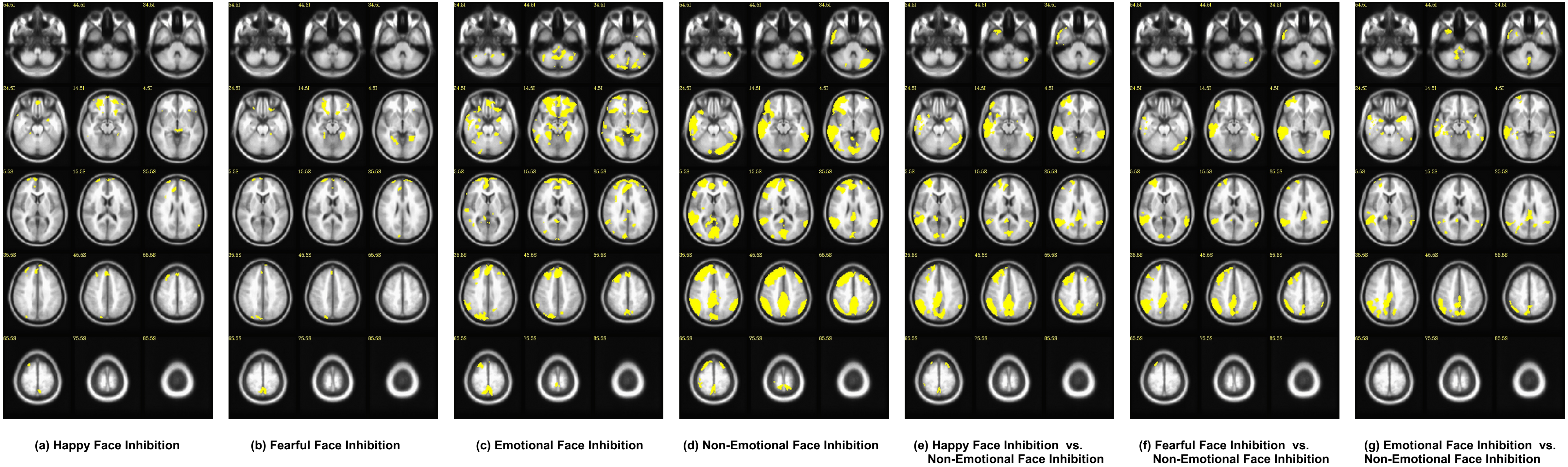


**Figure S-2.** Average effects of condition masks across all subjects at cluster-wise corrected significance of $p < .05$ for each condition respectively. (a) Happy face inhibition, (b) fearful face inhibition, (c) emotional face inhibition, (d) non-emotional face inhibition, (e) happy face inhibition vs. non-emotional face inhibition, (f) fearful face inhibition vs. non-emotional face inhibition, and (g) emotional face inhibition vs. non-emotional face inhibition.

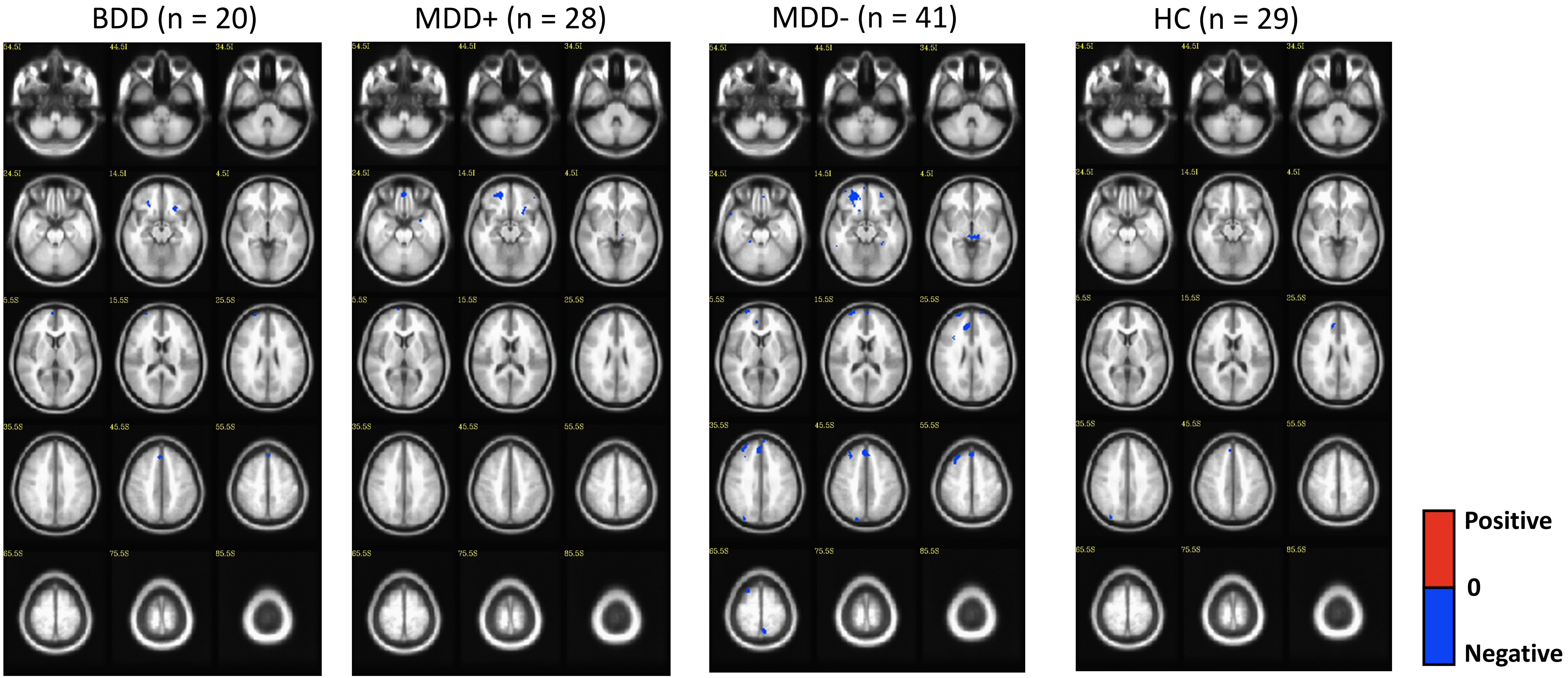


**Figure S-3.** Group activation during happy face inhibition. Significant activity within bipolar depressed (BD) group, major depressive disorder patients who are at high risk (MDD+) and low risk (MDD-) for developing BD, and healthy control (HC) group is shown at cluster-wise corrected significance of $p < .05$. A mask was created from average effect of condition across all subjects at cluster-wise corrected significance of $p < .05$, which was applied to group activation.

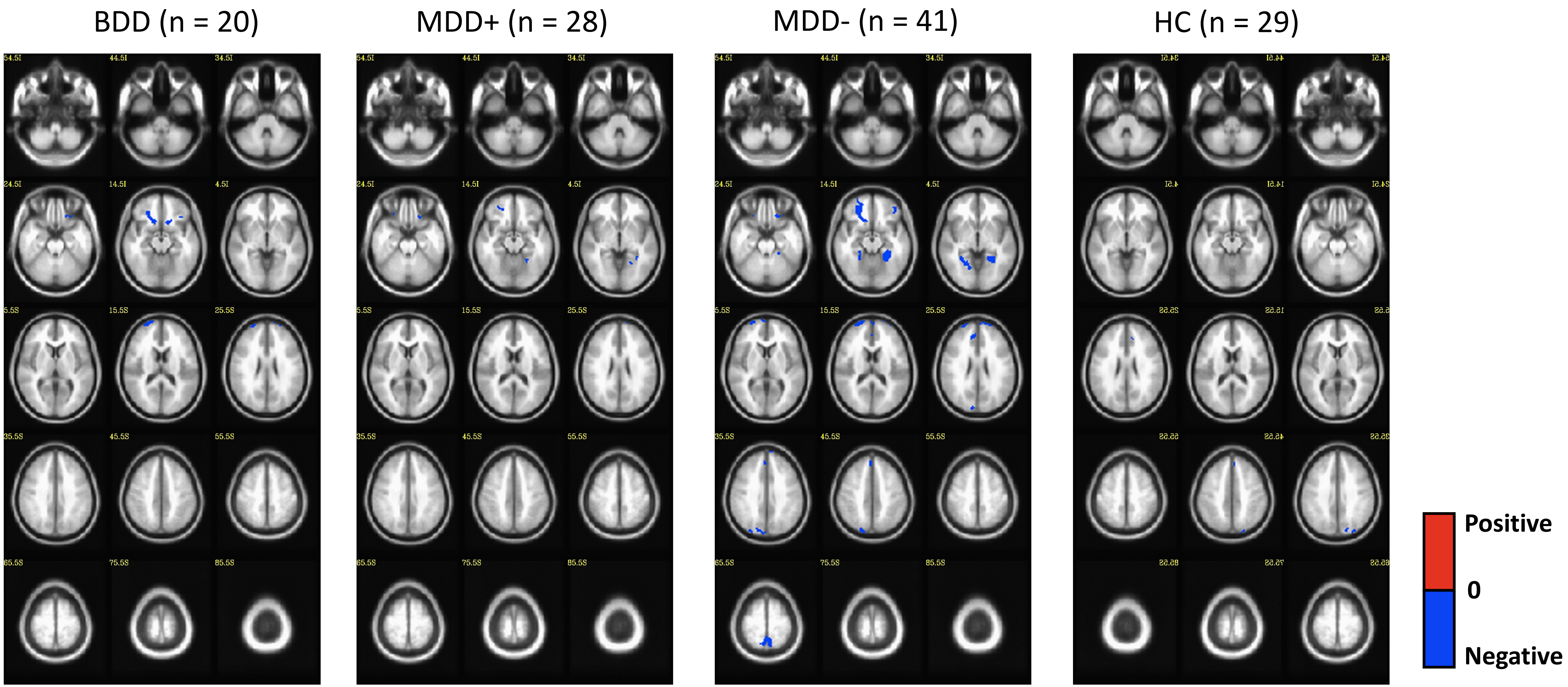


**Figure S-4.** Group activation during fearful face inhibition. Significant activity within bipolar depressed (BD) group, major depressive disorder patients who are at high risk (MDD+) and low risk (MDD-) for developing BD, and healthy control (HC) group is shown at cluster-wise corrected significance of $p < .05$. A mask was created from average effect of condition across all subjects at cluster-wise corrected significance of $p < .05$, which was applied to group activation.

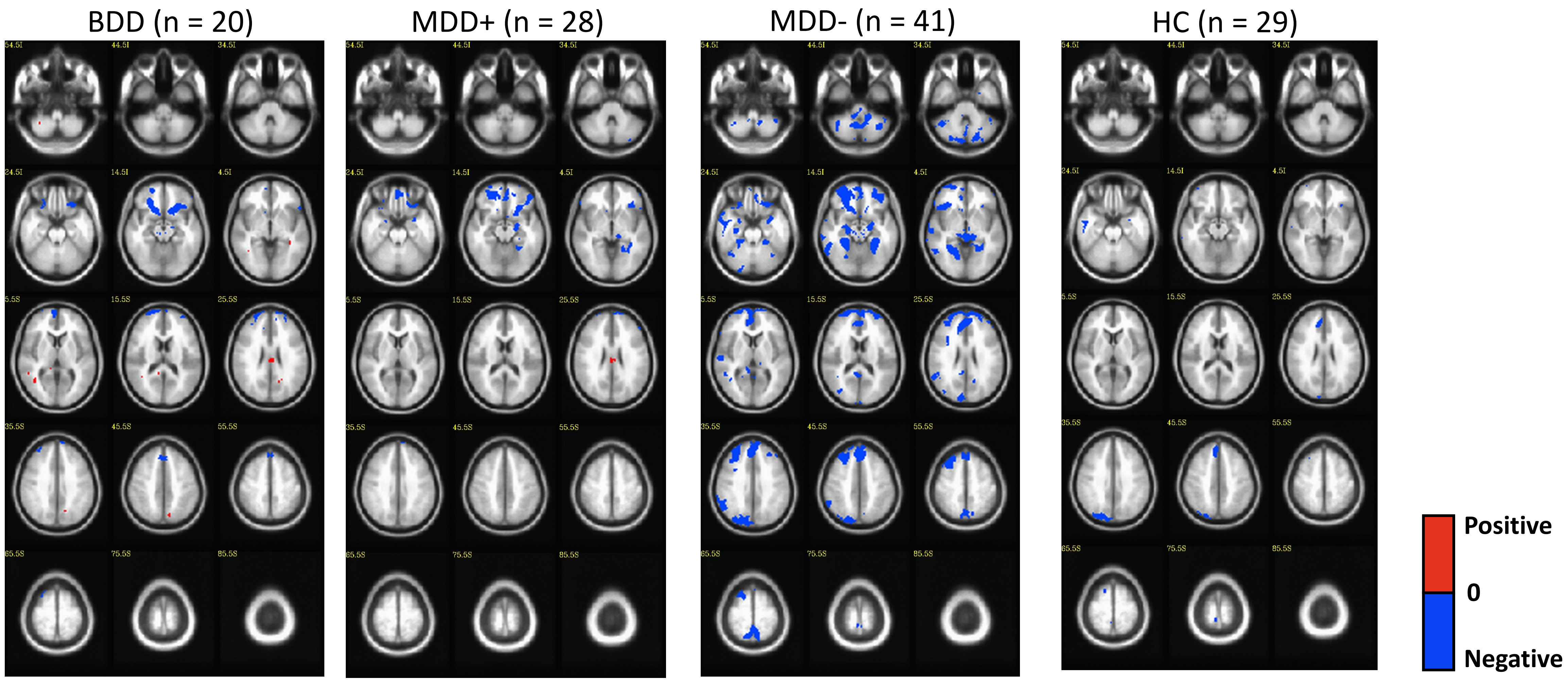


**Figure S-5.** Group activation during emotional face inhibition. Significant activity within bipolar depressed (BD) group, major depressive disorder patients who are at high risk (MDD+) and low risk (MDD-) for developing BD, and healthy control (HC) group is shown at cluster-wise corrected significance of $p < .05$. A mask was created from average effect of condition across all subjects at cluster-wise corrected significance of $p < .05$, which was applied to group activation.

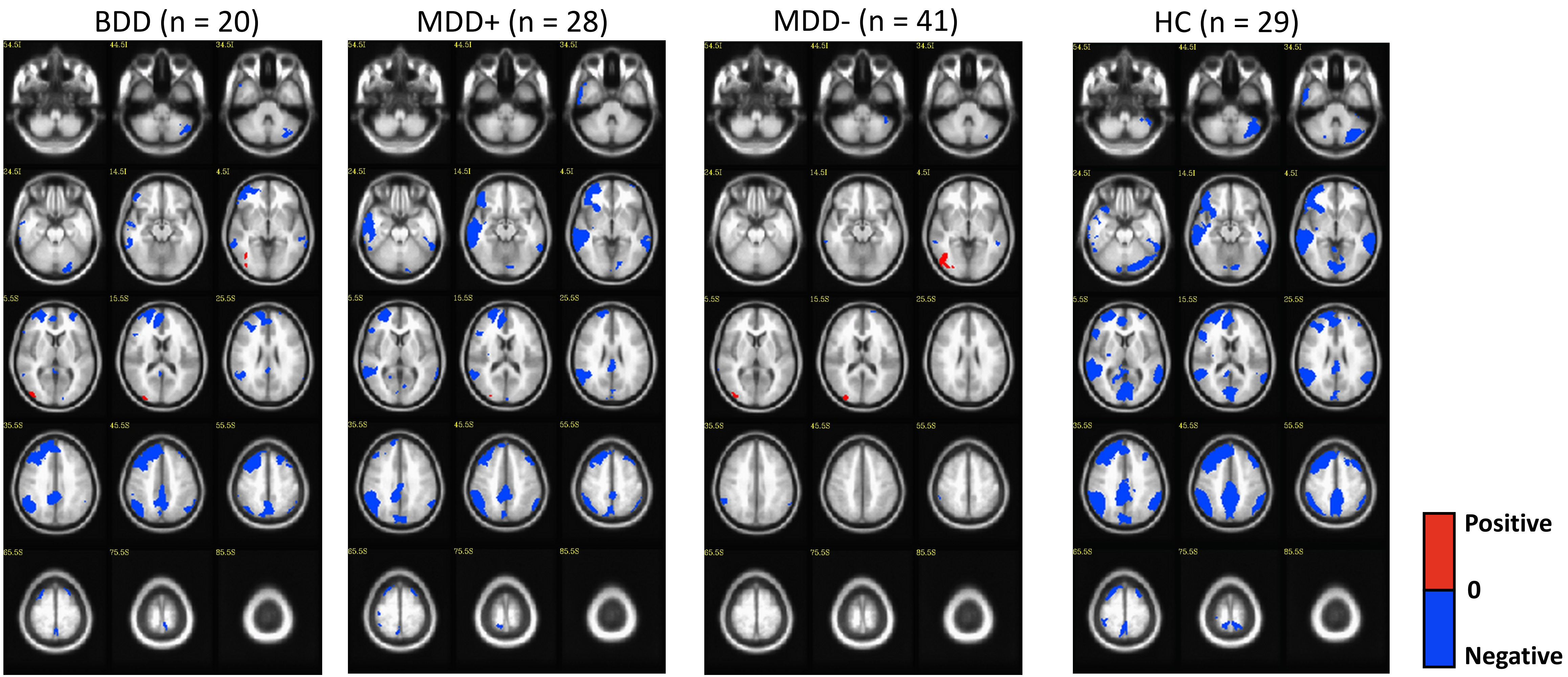


**Figure S-6.** Group activation during non-emotional face inhibition. Significant activity within bipolar depressed (BD) group, major depressive disorder patients who are at high risk (MDD+) and low risk (MDD-) for developing BD, and healthy control (HC) group is shown at cluster-wise corrected significance of $p < .05$. A mask was created from average effect of condition across all subjects at cluster-wise corrected significance of $p < .05$, which was applied to group activation.

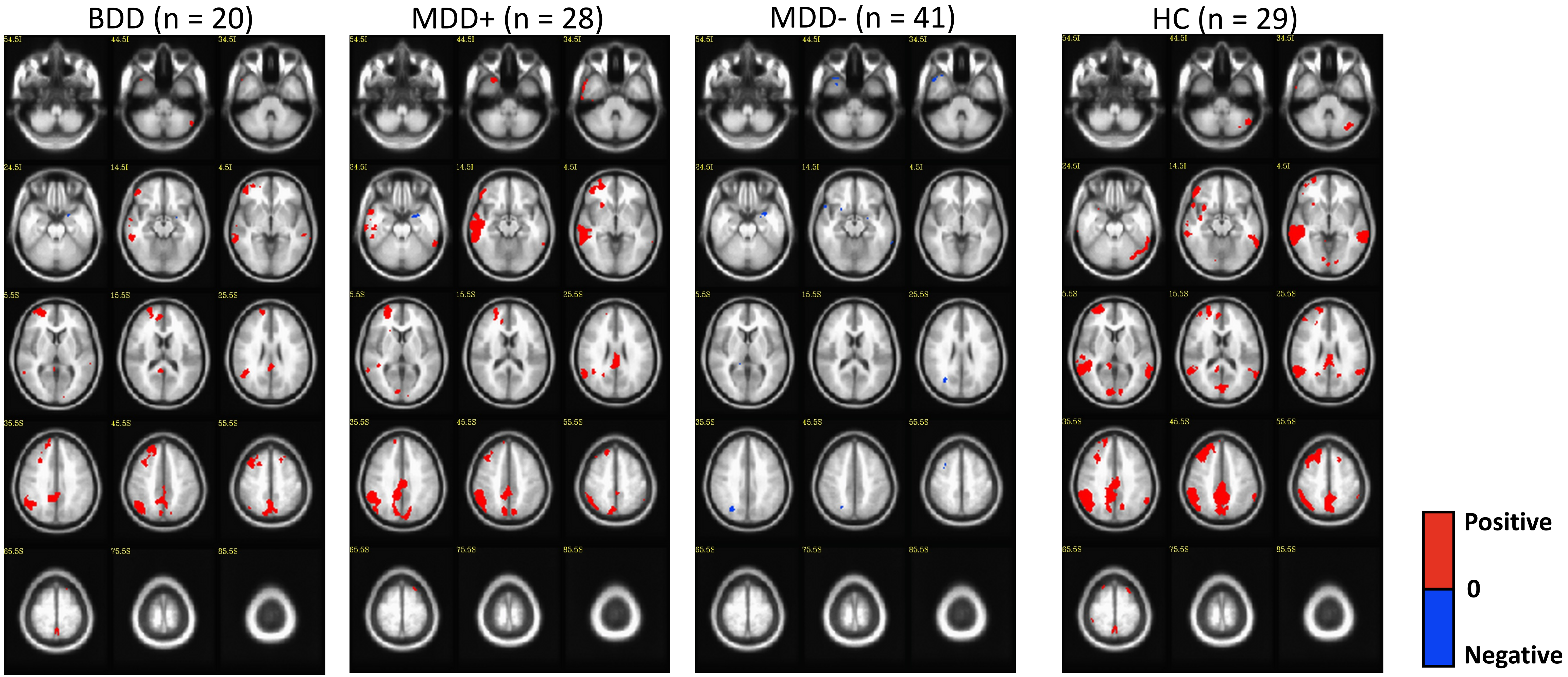


**Figure S-7.** Group activation during happy face inhibition vs. non-emotional face inhibition. Significant activity within bipolar depressed (BD) group, major depressive disorder patients who are at high risk (MDD+) and low risk (MDD-) for developing BD, and healthy control (HC) group is shown at cluster-wise corrected significance of $p < .05$. A mask was created from average effect of condition across all subjects at cluster-wise corrected significance of $p < .05$, which was applied to group activation.

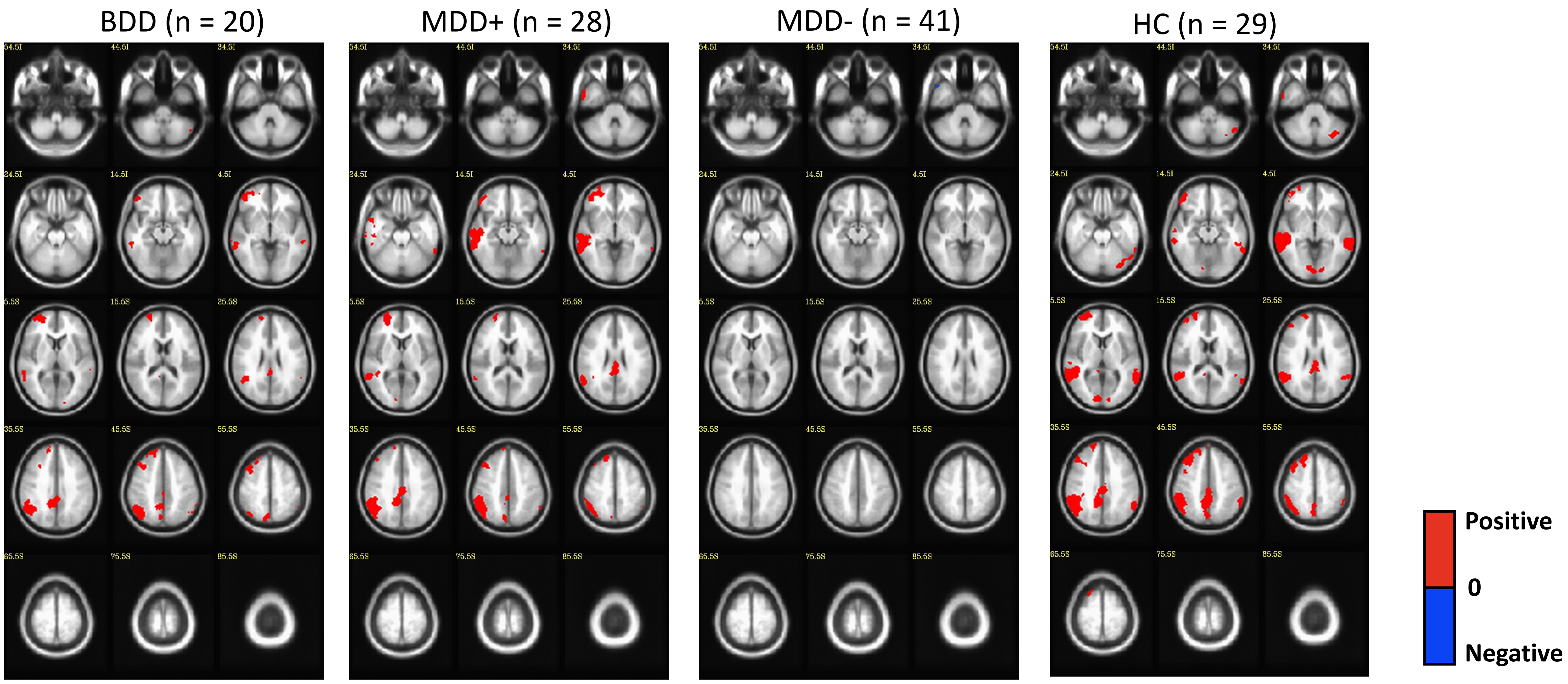


**Figure S-8.** Group activation during fearful face inhibition vs. non-emotional face inhibition. Significant activity within bipolar depressed (BD) group, major depressive disorder patients who are at high risk (MDD+) and low risk (MDD-) for developing BD, and healthy control (HC) group is shown at cluster-wise corrected significance of $p < .05$. A mask was created from average effect of condition across all subjects at cluster-wise corrected significance of $p < .05$, which was applied to group activation.

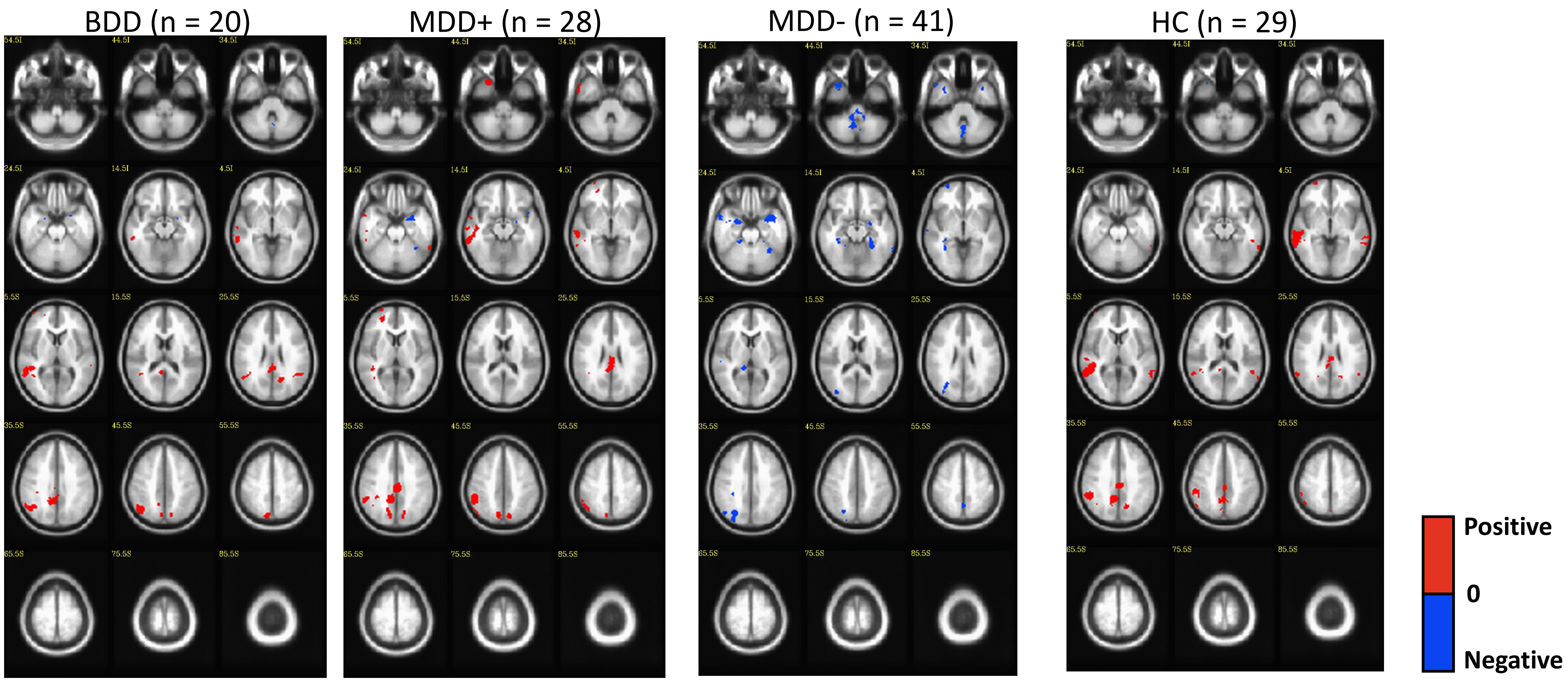


**Figure S-9.** Group activation during emotional face inhibition vs. non-emotional face inhibition. Significant activity within bipolar depressed (BD) group, major depressive disorder patients who are at high risk (MDD+) and low risk (MDD-) for developing BD, and healthy control (HC) group is shown at cluster-wise corrected significance of $p < .05$. A mask was created from average effect of condition across all subjects at cluster-wise corrected significance of $p < .05$, which was applied to group activation.

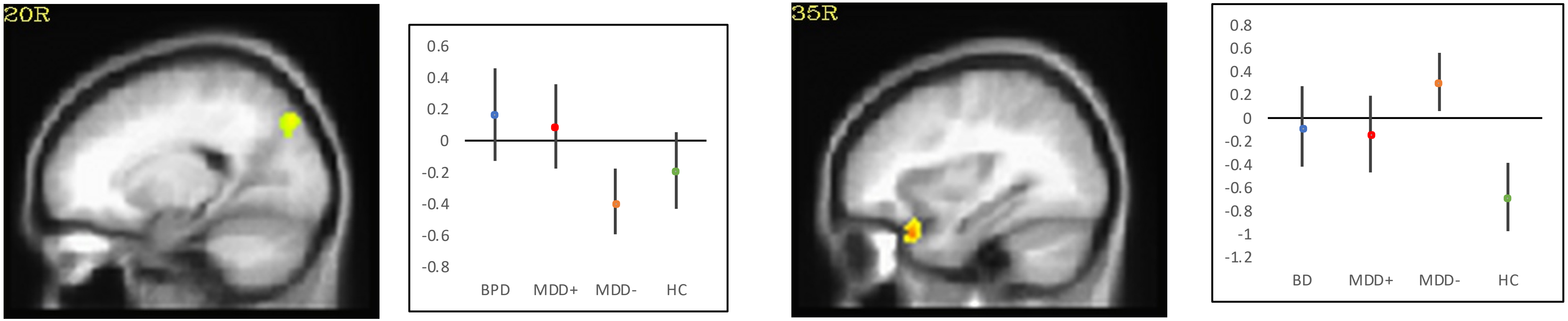


Right cuneus

Right medial temporal pole

(a) During emotional face inhibition

(b) During non-emotional face inhibition

**Figure S-10.** Group activation differences during emotional face inhibition (a) and during non-emotional face inhibition (b). The figure are shown at cluster size k $\geq$ 70 voxels for emotional face inhibition and K $\geq$ 57 voxels for non-emotional face inhibition, cluster-wise corrected significance of $p < .01$. Mean and 95% confidence intervals for average activity within main effect cluster are represented.

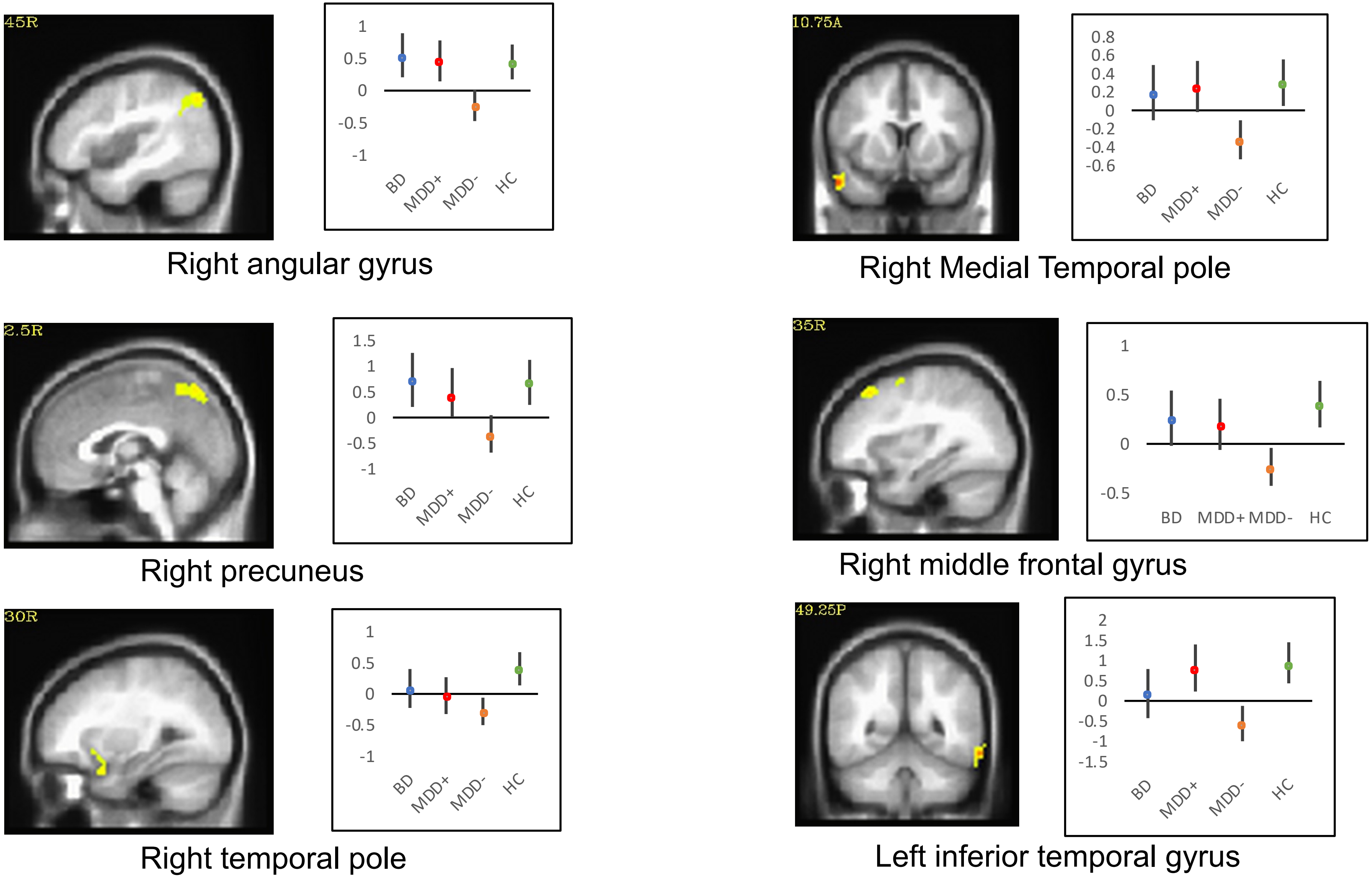


**Figure S-11.** Group activation differences during happy face inhibition vs. non-emotional face inhibition. The figure are shown at cluster size k $\geq$ 62 voxels, cluster-wise corrected significance of $p < .01$. Mean and 95% confidence intervals for average activity within main effect cluster are represented.

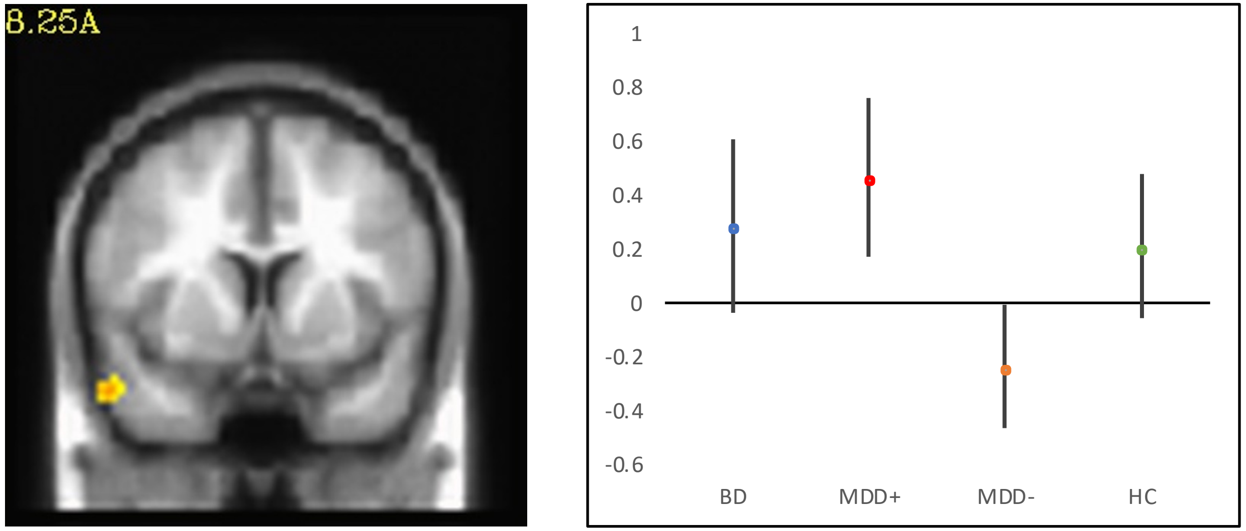


Right medial temporal pole

**Figure S-12.** Group activation differences during fearful face inhibition vs. non-emotional face inhibition. The figure is shown at cluster size k $\geq$ 42 voxels, cluster-wise corrected significance of $p < .01$. Mean and 95% confidence intervals for average activity within main effect cluster are represented.

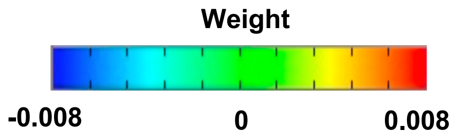


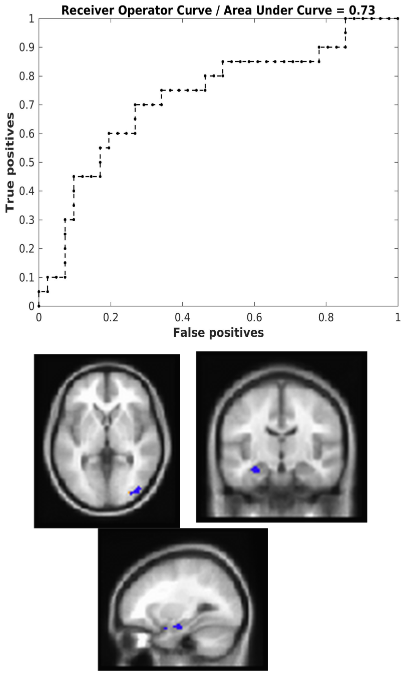


| Region of Interest | Weight |
|---|---|
| Right Middle Occipital Gyrus | -0.086051 |
| Left Hippocampus | -0.072868 |
| Right Cerebellum | 0.069277 |
| Right Precuneus | 0.037952 |
| Right Angular Gyrus | 0.037485 |

**(a) BDD vs. MDD-**

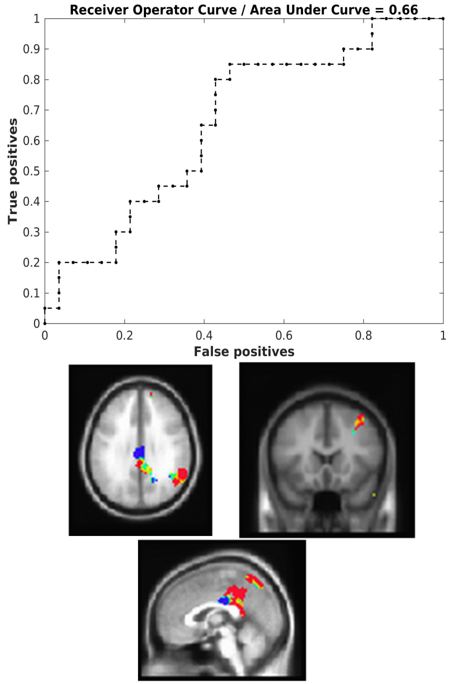


| Region of Interest | Weight |
|---|---|
| Right Inferior Temporal Gyrus | -0.057029 |
| Right Middle Cingulate Cortex | -0.035177 |
| Right Angular Gyrus | 0.032692 |
| Right Middle Temporal Gyrus | 0.026778 |
| Right Precuneus | 0.020676 |

**(b) BDD vs. MDD+**

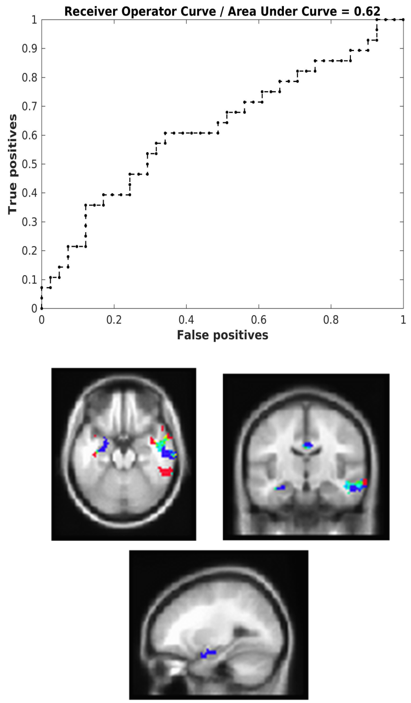


| Region of Interest | Weight |
|---|---|
| Right Angular Gyrus | 0.071290 |
| Left Hippocampus | -0.052000 |
| Right Fusiform Gyrus | 0.049272 |
| Left Temporal Pole | -0.042443 |
| Right Middle Temporal Gyrus | -0.039475 |

**(c) MDD+ vs. MDD-**

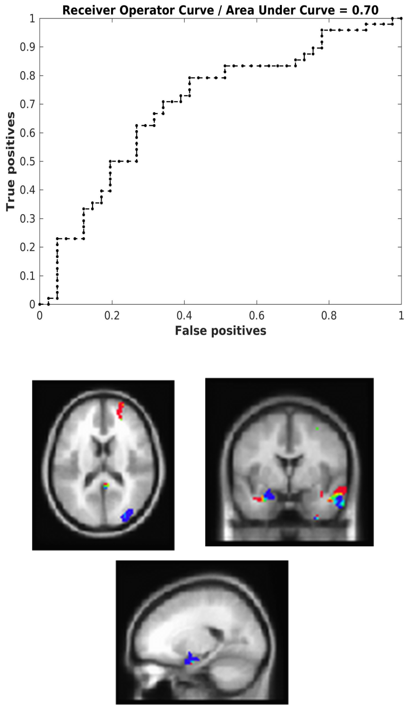


| Region of Interest | Weight |
|---|---|
| Right Superior Frontal Gyrus | 0.043852 |
| Right Inferior Frontal Gyrus | -0.041508 |
| Right Angular gyrus | 0.041382 |
| Right Temporal Pole | 0.039336 |
| Right Middle Frontal Gyrus | -0.022864 |

**(d) (BDD & MDD+) vs. MDD-**

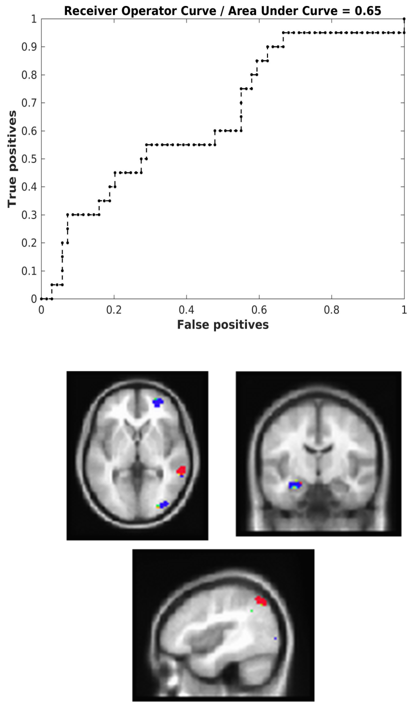


| Region of Interest | Weight |
|---|---|
| Left Temporal Pole | 0.069385 |
| Right Middle Occipital Gyrus | -0.048457 |
| Left Inferior Frontal Gyrus | 0.044981 |
| Right Precuneus | 0.042579 |
| Right Angular Gyrus | 0.041325 |

**(e) BDD vs. (MDD+ & MDD-)**

**Figure S-13.** Receiver Operating Characteristics (ROC) Curves with Area Under Curve (AUC) and color weighted discrimination maps of classification for happy face inhibition vs. non-emotional face inhibition generated by classifier using beta images masked by average effect of condition mask and top five most weighted brain regions for classification. (a) BDD vs. MDD- classification. Positive weights (red) represent the voxels contributing to classification as a BDD subject, while negative weights (blue) represent the voxels contributing to classification as an MDD- subject. (b) BDD vs. MDD+ classification. Positive class: BDD subject, negative class: MDD+ subject. (c) MDD+ vs. MDD- classification. Positive class: MDD+ subject, negative class: MDD- subject. (d) (BDD & MDD+) vs. MDD- classification. Positive class: (BDD & MDD+) subject, negative class: MDD- subject. (e) BDD vs.(MDD+ & MDD-) classification. Positive class: BDD subject, negative class: (MDD+ & MDD-) subject.

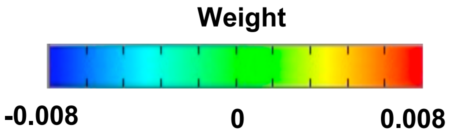


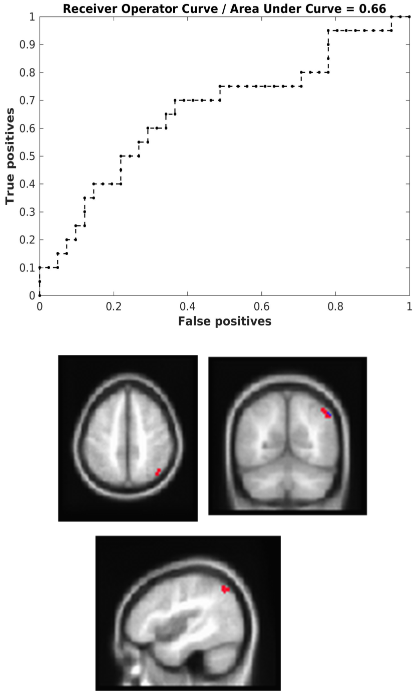


| Region of Interest | Weight |
|---|---|
| Right Angular gyrus | 0.236311 |
| Right Middle Temporal Gyrus | 0.100467 |

(a) BDD vs. MDD-

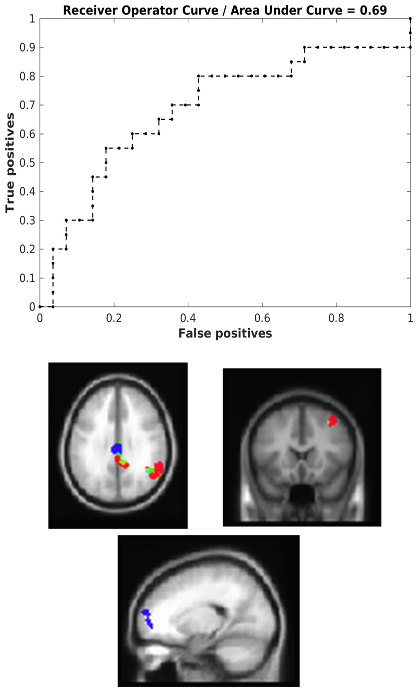


| Region of Interest | Weight |
|---|---|
| Right Middle Temporal Gyrus | 0.038772 |
| Right Middle Cingulate Cortex | -0.030462 |
| Right Angular Gyrus | 0.029015 |
| Right Superior Frontal Gyrus | -0.028476 |
| Right Middle Frontal Gyrus | 0.025576 |

(b) BDD vs. MDD+

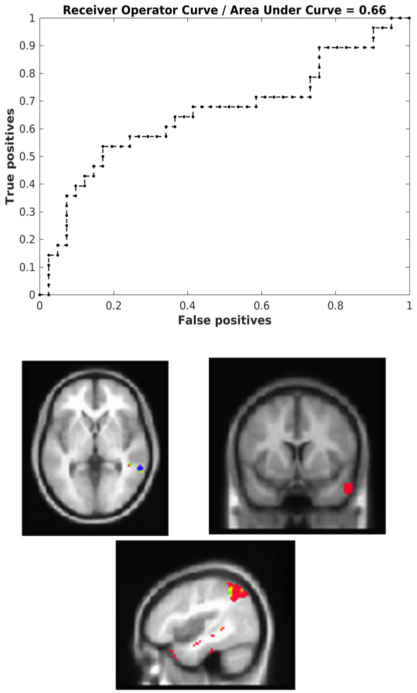


| Region of Interest | Weight |
|---|---|
| Right Angular Gyrus | 0.058589 |
| Right Inferior Temporal Gyrus | 0.053682 |
| Right Inferior Parietal Lobule | 0.044036 |
| Right Medial Temporal Pole | 0.041636 |
| Right Middle Temporal Gyrus | -0.041622 |

(c) MDD+ vs. MDD-

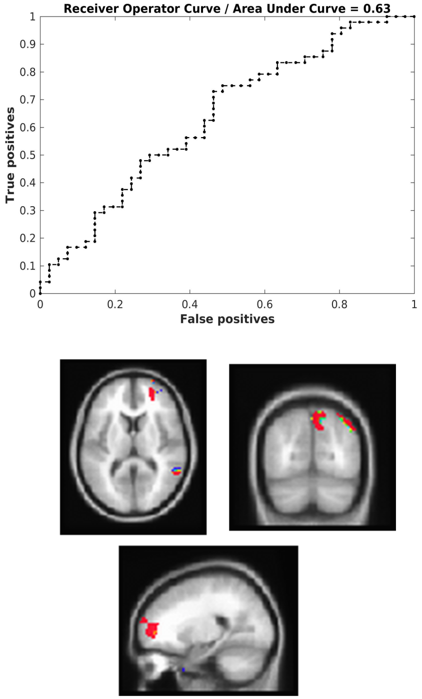


| Region of Interest | Weight |
|---|---|
| Right Superior Frontal Gyrus | 0.069788 |
| Right Middle Temporal Gyrus | -0.051324 |
| Right Inferior Temporal Gyrus | 0.050839 |
| Right Medial Temporal Pole | 0.045769 |
| Right Precuneus | 0.045053 |

(d) (BDD & MDD+) vs. MDD-

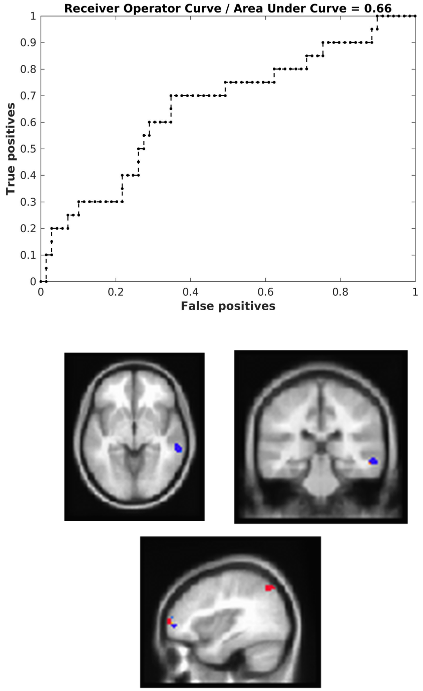


| Region of Interest | Weight |
|---|---|
| Right Middle Frontal Gyrus | 0.106570 |
| Right Inferior Temporal Gyrus | 0.096436 |
| Left Temporal Pole | 0.054781 |
| Right Angular Gyrus | 0.052806 |
| Right Middle Temporal Gyrus | -0.043510 |

(e) BDD vs. (MDD+ & MDD-)

**Figure S-14.** Receiver Operating Characteristics (ROC) Curves with Area Under Curve (AUC) and color weighted discrimination maps of classification for fearful face inhibition vs. non-emotional face inhibition generated by classifier using beta images masked by average effect of condition mask and top five most weighted brain regions for classification (a) BDD vs. MDD- classification. Positive weights (red) represent the voxels contributing to classification as a BDD subject, while negative weights (blue) represent the voxels contributing to classification as an MDD- subject. (b) BDD vs. MDD+ classification. Positive class: BDD subject, negative class: MDD+ subject. (c) MDD+ vs. MDD- classification. Positive class: MDD+ subject, negative class: MDD- subject. (d) (BDD & MDD+) vs. MDD- classification. Positive class: (BDD & MDD+) subject, negative class: MDD- subject. (e) BDD vs.(MDD+ & MDD-) classification. Positive class: BDD subject, negative class: (MDD+ & MDD-) subject.
